# Gut Commensal Bacteria-Derived Methionine is Required for Host Reproduction by Modulating RNA m6A Methylation of the Insulin Receptor

**DOI:** 10.1101/2024.08.20.608724

**Authors:** Qiuyuan Zhang, ZhuRong Deng, Xiaoxue Li, Jiao Qiao, Ziniu Li, Peipei Liu, Alfred M. Handler, Bruno Lemaitre, Weiwei Zheng, Hongyu Zhang

## Abstract

Gut commensal bacteria promote host reproduction by modulating metabolism and nutrition, yet the molecular mechanisms by which microbes modulate reproduction remain unclear. Here, we show that gut commensal bacteria promote host reproduction by providing amino-acid methionine, which controls the RNA m6A modification level of *insulin receptor* (*InR*) in the ovary of the invasive insect *Bactrocera dorsalis*. RNA m6A levels and S-adenosyl-methionine (SAM) titers in the ovaries were sharply reduced in antibiotic treated *B. dorsalis* compared with untreated insects, resulting in arrested ovarian development and decreased fecundity. The intestinal commensal bacteria *Enterobacter hormaechei* or the *E. hormaechei*-derived metabolite methionine restored the decreased RNA m6A level and the reproductive defects. Notably, knockdown of *METTL3* and *METTL14*, two genes encoding the RNA m6A methyltransferases, led to a decrease in the mRNA level of *InR* and underdevelopment of ovaries in *B. dorsalis,* and blocked the promoting effect of methionine on ovarian development and fecundity. Collectively, our study identifies an unrecognized role of RNA m6A methylation modification that underlies microbial control of host reproduction. Our findings further expand the functional landscape of m6A modification to include nutrient-dependent control of ovarian development and highlight the essential role of epigenetic regulation in microbe-host interactions.

## Introduction

Successful reproduction is essential for animals to maintain population stability and expansion. Gut microbiota has been recognized as a critical modulator of host physiology regulating the gut bacterial community homeostasis ^1, 2^, metabolism ^3^, behavior ^4^, and reproduction ^5^. Growing evidence has revealed that gut microbiota promotes host reproduction by providing essential amino acids and proteins ^6^. Perturbation on gut microbiota is related to reproductive defects in humans ^7^. In *Drosophila melanogaster*, the commensal bacteria *Acetobacter pomorum* and *Lactobacillus plantarum* synthesize essential amino acids that improve host fertility ^8^. However, little is known about the precise molecular mechanisms underlying the regulatory effect of gut microbiota on host reproduction.

Epigenetic regulation is increasingly recognized as a potent mechanism influencing host physiology^9^. N6-methyladenosine (m6A) is the most common and abundant methylation modification of eukaryotic RNA molecules ^10^. m6A methylation is a dynamic and reversible chemical modification in which methyltransferase acts to modify mRNA, thereby modulating the expression of genes at the post-transcriptional level ^11^. Notably, m6A modification has been found in various organisms, including plants ^12^, vertebrates such as humans ^13^, mice ^14, 15^, and zebrafish ^16^, and invertebrates such as *D. melanogaster* ^17, 18^. In insects, RNA m6A modification is essential for diverse biological processes, such as determination of sex ^17, 18^, resistance to insecticide ^19^, and development of embryo ^20^. The potential roles of m6A methylation modification in host reproduction, have yet to be fully described.

Owing to their high reproductive capacity, insects are the most diverse and largest animal group on the Earth. Among them, the oriental fruit fly, *B. dorsalis*, is an invasive insect that threatens agricultural industries by damaging over 350 species of vegetables and fruits worldwide ^21^. Previous studies revealed that *B. dorsalis* possesses a fairly stable gut microbial community ^1, 2, 22^ that affects many host traits such as resistance to heat stress ^23^, reproductive behavior ^24^, and recovery from radiation^25^. Thus, *B. dorsalis* is a suitable model organism for studying how gut microbiota affects host reproduction.

Here, we found that gut commensal *E. hormaechei*-derived methionine modulates RNA m6A modifications of the insulin receptor (*InR*) gene in the ovary of the host *B. dorsalis*, thereby controlling ovarian development. The findings provide insights into the underlying mechanism of microbe-host reproduction interplay at the epigenetic level.

## Results

### Gut commensal bacteria *E. hormaechei*-derived metabolite methionine is required for the fertility of *B. dorsalis* females

Ovarian development in *B. dorsalis* could be classified into five stages (I-V) according to the maturation degree (Supplementary Fig. 1a) ^26^. To determine whether the presence of gut bacteria in adult *B. dorsalis* females is required for ovary maturation, bacteria in *B. dorsalis* were eliminated by feeding the insect with streptomycin and penicillin antibiotics. Tissue dissection revealed that 100% of ovarian development was arrested in stages I and II in antibiotic treated (ABX) flies. Reinfection of germ-free *B. dorsalis* with total culturable bacteria rescued the ovarian developmental defect; 80% of ovaries were in stages IV and V, which is similar to those of the conventionally reared (CONV) groups (Supplementary Fig. 1b-d). To identify the microbiota species promoting ovarian development, mono-association experiments were further performed using six different bacterial strains isolated from the culturable gut microbiota of *B. dorsalis* (Supplementary Fig. 1e and Supplementary Table 1). Among the tested cultured bacteria, *E. hormaechei* had the most prominent ability to restore ovarian development of *B. dorsalis* (Fig. 1a, b, and Supplementary Fig. 1f, g). We further investigated the effect of *E. hormaechei* on fecundity by evaluating eggs laid capacity. The results showed that the ABX+EH group and the CONV group were not significantly different, while ABX treated animals were completely sterile (Supplementary Fig. 1h). In situ hybridization (FISH) revealed that *E. hormaechei* localizes to the midgut of *B. dorsalis* (Supplementary Fig. 1i). Additionally, we observed that feeding *B. dorsalis* with *E. hormaechei* fermentation broth also rescued host ovarian development (Supplementary Fig. 2a-c). These data indicate that *E. hormaechei* is related to ovarian development and fecundity in *B. dorsalis*.

**Figure 1:**
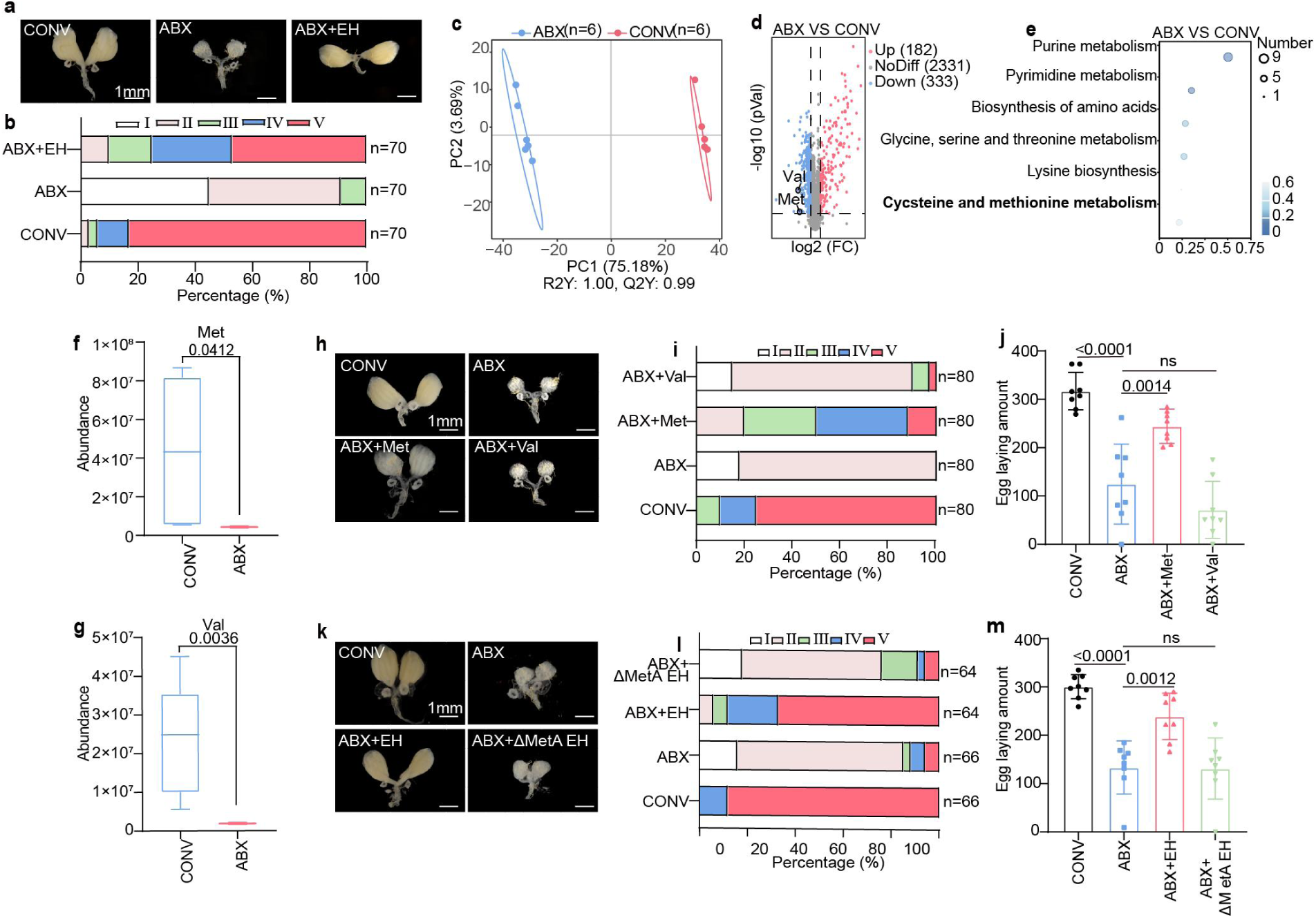
Methionine derived from gut commensal bacteria *E. hormaechei* is responsible for female reproduction in *B. dorsalis*. **a**, **b**, Morphology (a) and percentages of developmental stages (b) of ovaries in flies mono-associated with *E. hormaechei*. CONV, conventional flies; ABX, antibiotic treated flies; ABX+EH, ABX flies on a diet supplemented with *E. hormaechei.* **c**, Sparse PLS-DA score plot of hemolymph metabolic profiles of CONV and ABX flies. **d**, Volcano plots showing differentially regulated metabolites. Essential amino acids are highlighted (FDR<0.05 and |log2FoldChange|>1). **e**, Most enriched KEGG pathways downregulated in the hemolymph of ABX flies compared to CONV flies based on the differentially regulated amino acids (Fisher’s exact test). **f**, **g**, Abundance of Met (**f**) and Val (**g**) in *B. dorsalis* hemolymph in CONV and ABX flies. Data are from 3 independent biological replicates. **h-j**, Representative ovary morphology (**h**), percentages of developmental stages of ovaries (**i**), and total laid egg number during oviposition (**j**) ABX flies supplemented with Met (ABX+Met) or Val (ABX+Val). **k**-**m**, Representative ovary morphology (**k**), percentages of developmental stages of ovaries (**l**), and total number of eggs laid during oviposition (**m**) in ABX flies mono-associated with wild-type or mutant *E. hormaechei* strain (ΔMetA *E. hormaechei*, MetA deleted *E. hormaechei* mutant strain). Scale bar = 100 µm.

We speculated that *E. hormaechei* might produce a specific metabolite required for host ovarian development. To test this hypothesis, we performed untargeted metabolomics of host hemolymph and targeted metabolomics of *E. hormaechei* fermentation broth in MS medium. Untargeted metabolomics analysis of host hemolymph revealed significant differences in metabolite levels between ABX females and CONV groups as uncovered by principal component analysis (Fig. 1c). We found 515 metabolites whose levels differed between the two groups, with 333 metabolites downregulated in ABX females compared with CONV groups (Fig. 1d). KEGG enrichment analysis showed significant enrichment of the pathways associated with amino acid biosynthesis and metabolism, including cysteine and methionine metabolism, and lysine biosynthesis (Fig. 1e). Targeted metabolomics analysis of *E. hormaechei* revealed that this bacterium synthesized nearly all essential amino acids for the insect host except Lys (Supplementary Table 2). Integrated analysis of untargeted and targeted metabolomics revealed a significant reduction of the essential amino acids methionine (Met) and Valine (Val) in the hemolymph of ABX treated *B. dorsalis*, but no changes in other essential amino acids (Fig. 1d, f and g). Therefore, we examined how dietary supplementation with Met and Val would affect ovarian development of female *B. dorsalis*. Exogenous addition of Met significantly rescued ovarian developmental arrest in ABX treated animals, and about 50% of ovaries were at stage IV and stage V. In contrast, only 3% of ovaries in stage IV and V were significantly restored by supplementing Val (Fig. 1h, i). Addition of Met successfully restored egg-laying to wild-type levels in ABX insects (Fig. 1j), indicating that Met controls female reproduction in *B. dorsalis*. In contrast, Val did not restore the number of eggs laid.

To further determine the role of *E. hormaechei*-derived Met in the ovary development and fecundity of female *B. dorsalis*, we successfully constructed a knockout mutant (ΔMetA) disrupted in the Met synthesis pathway (Supplementary Fig. 3a-c). The ΔMetA *E. hormaechei* was able to colonize the host gut (Supplementary Fig. 3d) and did not show any growth defect in LB medium (Supplementary Fig. 3f). We further found that the content of Met in the *B. dorsalis* hemolymph decreased by 1.7-fold in ABX females compared with the CONV group (Supplementary Fig. 3e). Supplementation with wild type *E. hormaechei* but not with the ΔMetA *E. hormaechei* strain restored the Met content (Supplementary Fig. 3e), the ovarian phenotypic defects (Fig. 1k, l), and the laid egg number (Fig. 1m), with no significant difference compared with the CONV group. Taken together, the above results suggested that the gut bacteria-derived essential amino acid Met is required for female reproduction in *B. dorsalis*.

### Met is essential for RNA m6A methylation in the ovary of *B. dorsalis*

RNA m6A methylation regulates many physiological processes in eukaryotes ^12, 15, 27^. Thus, we explored whether the microbiota-derived metabolite Met could affect ovarian development by affecting m6A methylation. We first performed dot blot assays and quantified the m6A levels in the ovary of *B. dorsalis* using m6A methylation-specific antibodies. No obvious m6A signal was detected in ABX insects compared with CONV flies (Fig. 2a, b). Notably, feeding *B. dorsalis* with Met significantly increased the RNA m6A methylation level in ABX treated insects (Fig. 2a). Strikingly, supplementation with wild-type but not the -ΔMetA *E. hormaechei* fully restored m6A methylation levels in ABX treated insects (Fig. 2b). Quantification of total m6A levels in *B. dorsalis* hemolymph by ELISA showed similar results to those obtained in dot blot assays: dietary supplementation of ABX flies with Met (Fig. 2c) or WT *E. hormaechei* (Fig. 2d) increased the m6A level by 1.8 and 2.2 folds, respectively. Similarly, the titer of SAM, the donor of RNA m6A modification ^28, 29^, significantly decreased in the hemolymph of ABX treated compared to CONV raised *B. dorsalis*, while it was restored to wild-type level by supplementation of wild type *E. hormaechei* strains but not the ΔMetA *E. hormaechei* strains (Fig. 2e, f). These results were also confirmed by MeRIP-seq data (Supplementary Fig. 4). Principal component analysis (PCA) of m6A peaks by m6A sequencing clearly separated the methylation peaks of CONV and ABX in the ovary, but m6A marks of ABX flies fed with *E. hormaechei* (ABX+EH) were clustered together with the methylation peaks of CONV individuals (Supplementary Fig. 4).

**Figure 2:**
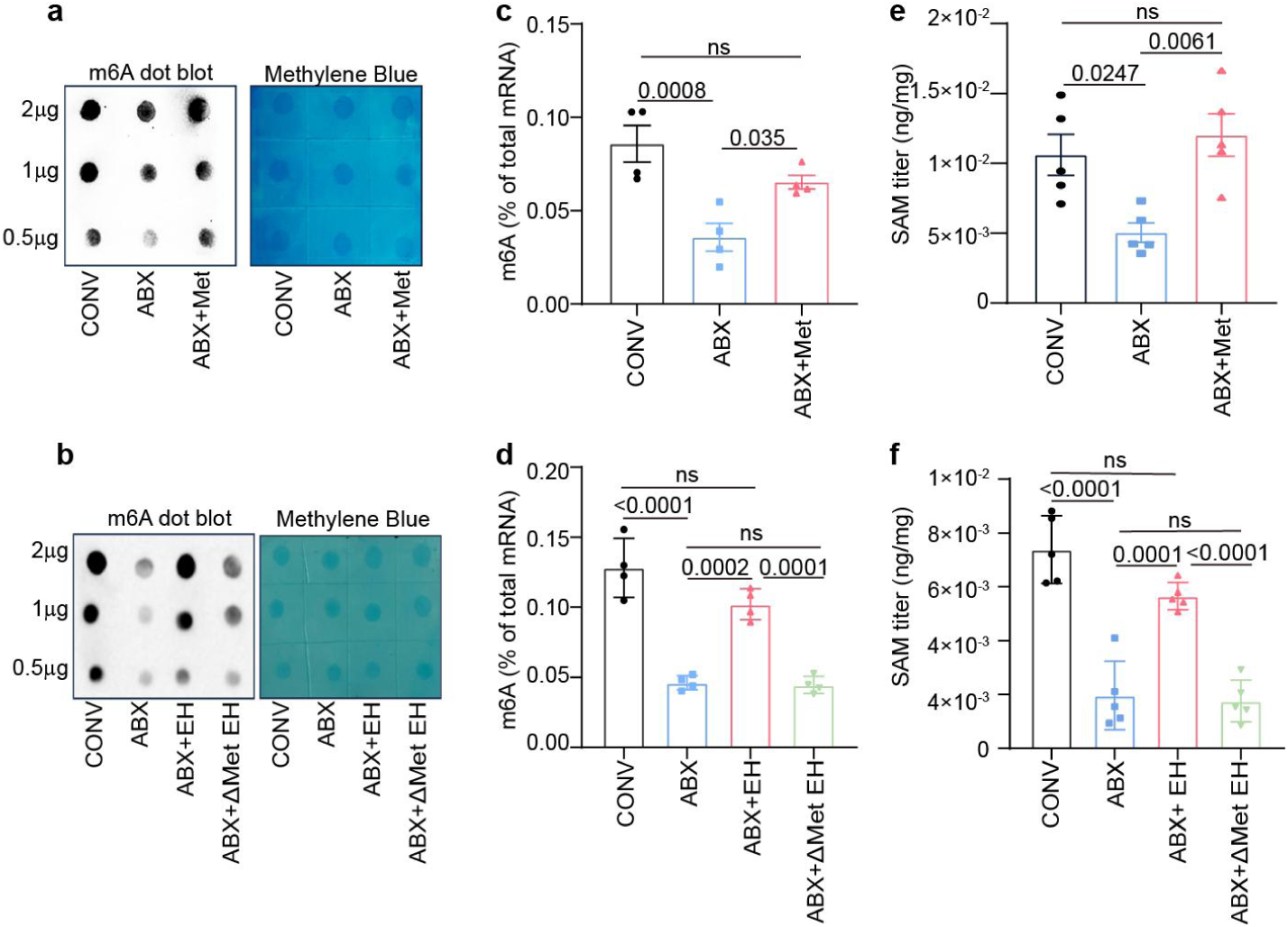
Methionine is essential for RNA m6A methylation levels in the ovary of *B. dorsalis*. **a**,**b**, RNA dot blot analysis on m6A level in the CONV, ABX, and ABX+Met groups (a) as well as in the CONV, ABX, ABX+ *E. hormaechei*, and ABX+ ΔMetA *E. hormaechei* groups (b). RNA samples were serially diluted and equal amounts of 2, 1, and 0.5 μg were loaded for each treatment group. Methylene blue staining was carried out to serve as a loading control. **c**,**d**, ELISA-measured mRNA m6A levels in the CONV, ABX, and ABX+Met (c) as well as in the CONV, ABX, ABX+WT *E. hormaechei*, and ABX+ ΔMetA *E. hormaechei* groups (d). **e**, **f**, Titer of SAM in the hemolymph of *B. dorsalis* in the CONV, ABX, and ABX+Met (e) as well as in the CONV, ABX, ABX+WT *E. hormaechei*, and ABX+ ΔMetA *E. hormaechei* groups (f). The number of independent biological replicates was four for **c** and **d**, while five for **e** and **f**.

METTL3 and METTL14 are two core enzymes of the RNA m6A methylation pathway that act as ‘writers’. Writer enzymes cooperate to transfer a methyl group from SAM to the N6 amino group of an adenosine (A) base in RNA to convert it to m6A ^30, 31^. We observed that the METTL3 and METTL14 protein levels decreased in the ovary of *B. dorsalis* of ABX treated females compared to CONV animals (Supplementary Fig. 5a, b). The expression levels of *METTL3* and *METTL14* were restored when flies were fed with Met or wild *E. hormaechei* but not ΔMetA *E. hormaechei* strains (Supplementary Fig. 5a, b). Taken together, these data indicated that the presence of Met provided by the microbiota is critical for m6A RNA methylation levels in the ovary of *B. dorsalis*, raising the hypothesis that defects in this pathway might be the cause of ovarian development arrest in germ free animals.

### RNA m6A methylation modifications are required for female reproduction of *B. dorsalis*

To investigate whether m6A methylation modifications are necessary for female reproduction, we first explored the spatiotemporal pattern of m6A modification in *B. dorsalis.* Over time, m6A was enriched during embryogenesis peaking at 12h, but dropped at the larval stage to remain low in the pupa, and reached its highest level in female adults (Supplementary Fig. 6a). In the adult insect, m6A was mainly enriched in the ovaries (Supplementary Fig. 6b). Furthermore, the mRNA expression levels of METTL3 and METTL14 were evaluated by RT-qPCR, and their developmental patterns were consistent with that of m6A (Supplementary Fig. 6a, b). *METTL3* and *METTL14* exhibited 5.58 and 5.60 times higher expression levels in female adults compared with male adults (Supplementary Fig. 6a). Furthermore, both genes were primarily enriched in the ovaries of females (Supplementary Fig. 6b), suggesting their involvement in female reproduction.

To further probe the role of m6A methylation modifications in *B. dorsalis* ovary development, RNA interference (RNAi) mediated knockdown of *METTL3* and *METTL14* was performed in female adults. Both the transcript (Supplementary Fig. 7a, b) and protein (Fig. 3a, b) levels of METTL3 and METTL14 were significantly reduced 48 h after injection of dsMETTL3 and dsMETTL14 in the body cavity. Dot blot assays indicated that knockdown of *METTL3* and *METTL14* reduced the total RNA m6A methylation level by 59% and 63% in treated flies compared with dsEGFP-injected females (Fig. 3c, d). Morphological analysis revealed that 70% of dsMETTL3-injected females and 80% of dsMETTL14-injected individuals displayed underdeveloped ovaries (Fig. 3e, f). Additionally, after *METTL3* or *METTL14* knockdown, the amount of egg-laying was significantly reduced by 5 folds or 2 folds, compared with the control group (Fig. 3g). Collectively, the above results suggested that m6A methylation modifications are necessary for ovarian development and fecundity.

**Figure 3:**
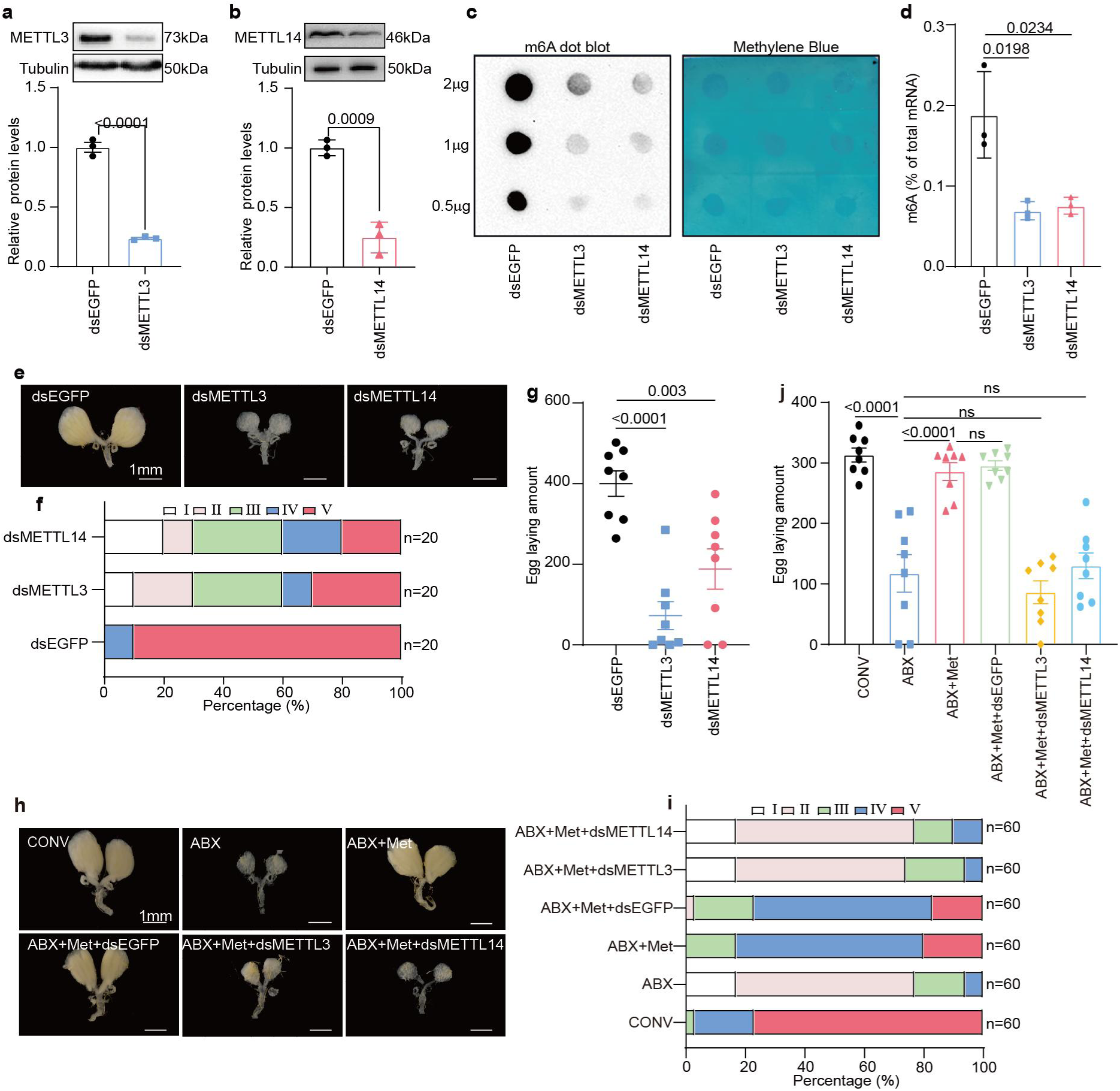
RNA m6A methylation modifications are required for female reproduction in *B. dorsalis*. **a**, **b**, RNAi-mediated knockdown of *METTL3* (a) and *METTL14* (b) was detected at the protein level. Tubulin served as a loading control. The Protein level was quantified by gray value and normalized to Tubulin. **c**, RNA dot blot analysis of m6A levels after insects were injected with dsEGFP, dsMETTL3, and dsMETTL14. Methylene blue staining was used as a loading control. **d**, ELISA-quantified mRNA m6A level in different groups injected with dsEGFP, dsMETTL3, or dsMETTL14. **e**-**g,** Effect of knockdown of *METTL3* or *METTL14* gene on ovarian development (**e**, **f**) and egg laying amounts (**g**) of *B. dorsalis*. **h-j**, Representative ovarian morphology (**h**) and percentages of developmental grades of ovaries (**i**), and total number of eggs laid during oviposition (**j**) after a diet supplemented with methionine in dsMETTL3-injected or dsMETTL14-injected females. The number of independent biological replicates was three for **a**, **b**, and **e**, while eight for **g** and **j**.

We further evaluated whether Met promotes ovarian development through m6A methylation modification in *B. dorsalis*. Our results showed that after knockdown of *METTL3* or *METTL14*, supplementation with Met failed to rescue ovarian development arrest (Fig. 3h, i) and reduced egg laying (Fig. 3j) in ABX treated animals. The METTL3 or METTL14 knock-down groups showed significantly lower percentages of stage IV and stage V ovaries (6% in ABX-Met-dsMETTL3 or 10% in ABX-Met-dsMETTL14) compared with the control or dsEGFP-injected groups (80% in ABX-Met-dsEGFP) at 10 days post-eclosion (Fig. 3h, i). Taken together, we confirmed that Met promotes ovarian development and fecundity by controlling RNA m6A methylation levels.

### *InR* is a target gene of m6A methylation controlling ovarian development

To identify potential target genes of m6A methylation modification that might regulate ovarian development, we conducted an integrated analysis on the RNA-seq and MeRIP-seq data of ovaries. We identified 43 genes whose hypomethylation down-regulated mRNA levels in ABX-treated animals compared to CONV or ABX+EH animals (Fig. 4a and Supplementary Table 3). Interestingly, KEGG pathway enrichment analysis of these genes showed enrichment in the FOXO and mTOR signaling pathways, both of which are associated with nutrient signaling and the insulin receptor gene *InR* was overlapped between both pathways (Fig. 4b). Notably, the insulin receptor signaling pathway was also enriched based on GO enrichment analysis (Fig. 4c). This raised the hypothesis that m6A methylation modification of mRNAs involved in nutritional metabolic pathways might play an important role in ovary development in *B. dorsalis*. Therefore, we specifically focused on genes of the insulin pathway. Interestingly, RNAseq revealed that *InR* expression was down-regulated in ABX compared to CONV flies (Fig. 4d), or ABX treated flies supplemented with *E. hormaechei* or fed with Met (Fig. 4e). RT-qPCR (Fig. 4f) and RNA immunoprecipitation qPCR (m6A-IP-qPCR) (Fig. 4g) validated the omics results showing that *InR* transcript levels and m6A levels of *InR* mRNA were reduced. We further found that Met rescued the reduction of mRNA (Fig. 4h) and mRNA m6A levels (Fig. 4i) of *InR* transcripts in ABX flies. The m6A-qPCR also showed that *InR* mRNA is a direct target of the m6A modification (Fig. 4i).

**Figure 4:**
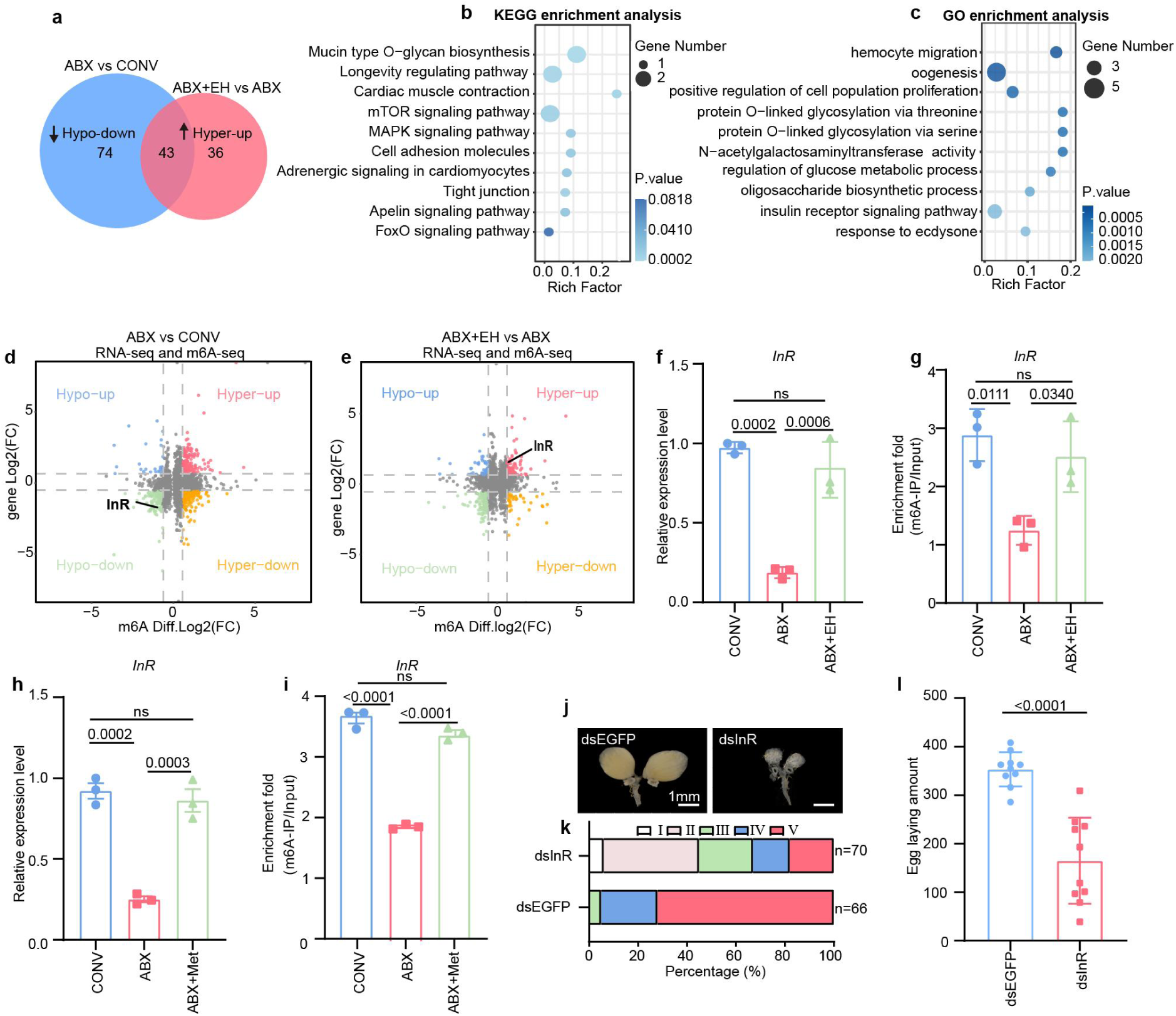
MeRIP-seq and RNA-seq analyses identify *InR* as a target gene controlling ovarian development in *B. dorsalis*. **a**, **b**, **c**, Venn diagram (a), KEGG enrichment analysis (b), and GO enrichment analysis (c) combines the hypomethylated-down mRNAs in the ABX group compared with CONV, and the hypermethylated-up mRNAs in the compensated *E. hormaechei* group compared with ABX group. **d**, **e**, Quadrant diagram representing combined analysis of RNA-seq and MeRIP-seq in the ABX vs. CONV (d) and ABX+EH vs. ABX groups (e). **f**, **g**, mRNA expression (f) and RNA m6A immunoprecipitation with quantitative PCR (g) of *InR* in the ovary of CONV, ABX, or ABX+EH flies. **h**, **i**, mRNA expression (h) and RNA m6A immunoprecipitation with quantitative PCR (i) of *InR* in the ovary of CONV, ABX, or ABX+Met flies. **j-l**, Effect of knockdown of *InR* on ovarian development, including representative ovarian morphology (**j**) and percentages of developmental stages of ovaries (**k**), and total number of eggs laid during oviposition (**l**). The number of independent biological replicates was three for **f**-**i**, while eight for **l**.

To elucidate whether the *InR* gene is essential for the reproduction of female *B. dorsalis*, RNAi-mediated knockdown of *InR* was performed by injecting dsRNA in females (Supplementary Fig. 8a). Tissue dissection revealed that *InR* knockdown led to underdeveloped ovaries (Fig. 4j, k), which is similar to those observed in antibiotic treated animals or dsMETTL3 or dsMETTL14 treated groups (Fig. 3e, f). Additionally, *InR* knockdown also reduced the laid egg number per female (Fig. 4l). Considering that the insulin signaling pathway controls female reproduction by interplay with 20-hydroxyecdysone (20E) during vitellogenesis in Diptera (Smykal and Raikhel, 2015), and the response to ecdysone pathway was also enriched based on GO enrichment analysis (Fig. 4c), we assayed 20E titer and the expression profiles of genes associated with ecdysone biosynthesis and 20E response genes such as vitellogenin (Vg), whose expression is activated by 20E, using RT-qPCR. The results showed that dsInR-injected females had significantly lower 20E titers than dsEGFP-injected individuals (Supplementary Fig. 8b), and ecdysone biosynthesis (*Phantom*, *Disembodied*, *Shroud*, *Neverland*, *Spook, Shadow*, *Shade*), 20E response (*E75*, *HR3*, *EcR*, *USP*, *E74*, *FTZ-f1*), and vitellogenin (Vg) were dramatically down-regulated as measured by RT-qPCR (Supplementary Fig. 8c). Taken together, these results support the conclusion that *InR* is essential for the reproduction of female *B. dorsalis*.

### Methylation sites in the 5′UTR of *InR* are identified in m6A-regulated *InR*

Our results demonstrated that *InR* mRNA can be m6A methylated (Fig. 4d, e, g) and MeRIP-sequencing data revealed that m6A was significantly enriched in its 5′UTR regions (Fig. 5a). Hence, we supposed that the expression of *InR* is possibly regulated by m6A modification. To verify this, *InR* was analyzed in the ovary of females upon knockdown of METTL3 and METTL14. The mRNA expression of *InR* dramatically declined by about 3 folds in both treated groups, compared to dsEGFP (Fig. 5b). Consistently, m6A-RIP-qPCR confirmed that the m6A modification level of *InR* mRNA decreased by about 7 folds and 5 folds in dsMETTL3-injected and METTL14-injected individuals, respectively (Fig. 5c). Further, the mRNA half-life of *InR* was significantly reduced in METTL3 or METTL14-knockdown embryonic cells of *B. dorsalis* compared to control cells (Fig. 5d). We then used the forecasting tool SNAMP to predict the m6A site of *InR* mRNA. Two putative m6A methylation sites were identified with high confidence on the 5’UTR region at 156 (-A156) and 217 (-A217) upstream of the start codon, which is consistent with our MeRIP-sequencing results (Fig. 5e, f). Subsequently, we mutated these two m6A sites by changing adenosine (A) to guanine (G) (-A156-G and -A217-G) and amplified the correct sequences into a vector with a luciferase reporter gene and minimal promoter (Fig. 5g). Dual-luciferase reporter assays were performed in embryonic cells of *B. dorsalis* to monitor the impact of these two m6A modification sites on mRNA stability *in vitro*. Following co-transfection of gene fusion recombinant plasmids into *B. dorsalis* cells, the luciferase activity of 5′UTR-InR2 reporter genes was significantly decreased when the m6A site -A156 was mutated, but there was no significant difference when the -A217 site was mutated (Fig. 5h). Together, these results indicated that the m6A site -A156 in the 5′UTR is responsible for mRNA stability of the *InR* gene *in vitro*, confirming that this gene is post-transcriptionally regulated by the m6A modification machinery.

**Figure 5:**
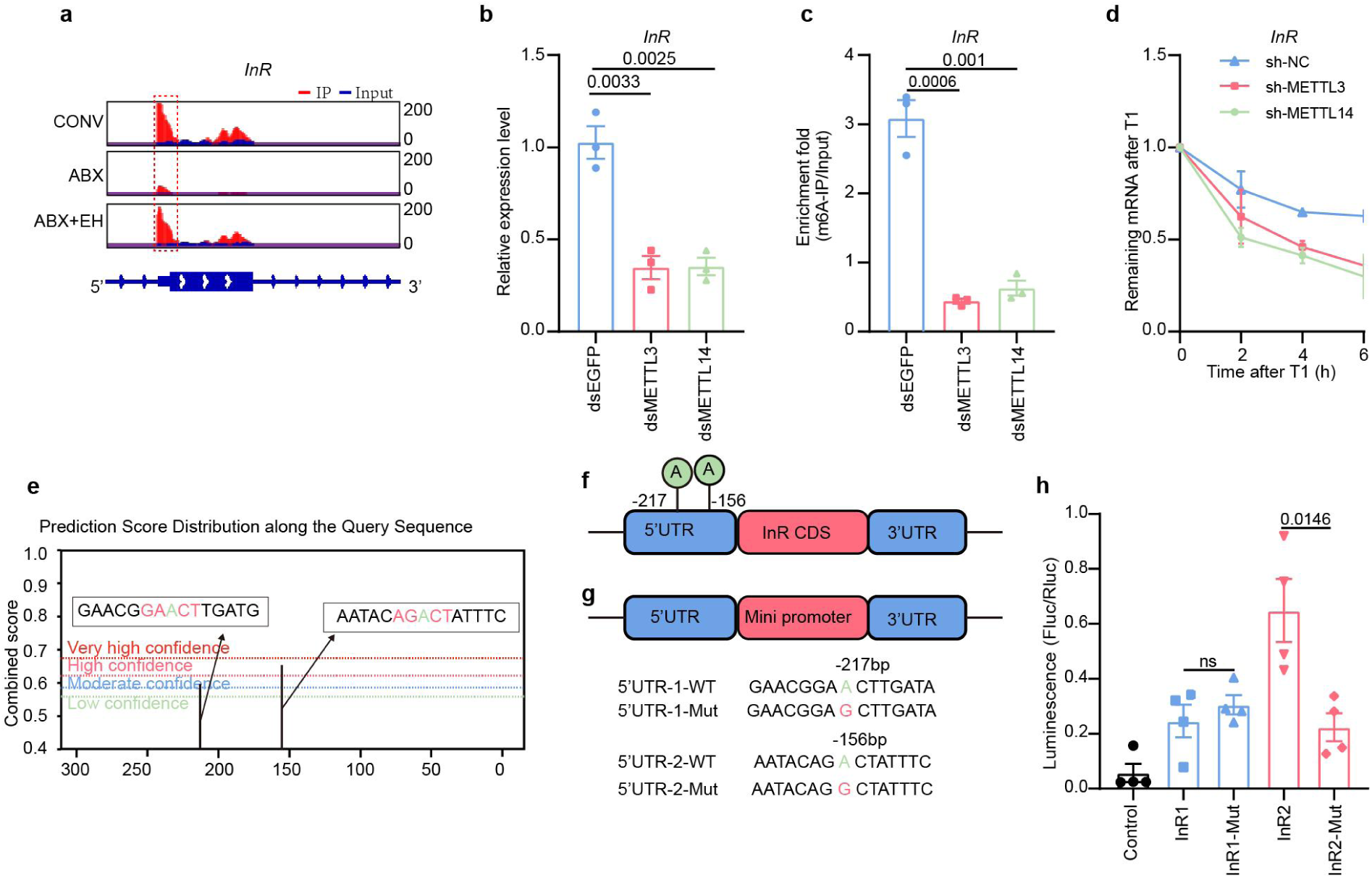
Methylation sites in the 5′UTR of *InR* are identified in m6A-regulated *InR.* **a**, IGV tracks showing m6A peaks in the 5′UTR of *InR* mRNA. **b**,**c**, mRNA expression (b) and RNA m6A immunoprecipitation with RT-qPCR (c) of *InR* in female flies injected with dsEGFP, dsMETTL3, or dsMETTL14. **d**, The RNA lifetimes of *InR* were checked in sh-NC, shMETTL14, and shMETTL14 in *B. dorsalis* cells after Act-D treatment for inhibition of transcription. **e**, Bioinformatic prediction of m6A sites of *InR*. **f**, Schematic of m6A motif positions within *InR* mRNA. **g**, Construction of a double luciferase reporter vector carrying the A156 and A217 mutations in the 5′UTR consensus sequence of *InR* (A217-G, A156-G). **h**, Luciferase reporter assays of candidate m6A modification sites (A156 and A217 in the 5’UTR of *InR*) *in vitro*. The number of independent biological replicates was three for **a-c**, while four for **d, g.**

## Discussion

This study revealed that gut microbiota is critical for host reproduction in the insect *B. dorsalis.* Gut bacteria, notably *E. hormaechei*, promotes ovarian development by producing an essential amino acid methionine. Deficiency in methionine particulary affects the m6A methylation pathway that regulates the stability of many transcripts required for ovarian development. Among them, we show that the mRNA stability of the insulin receptor gene *InR* depends on m6A methylation (Fig. 6).

**Figure 6:**
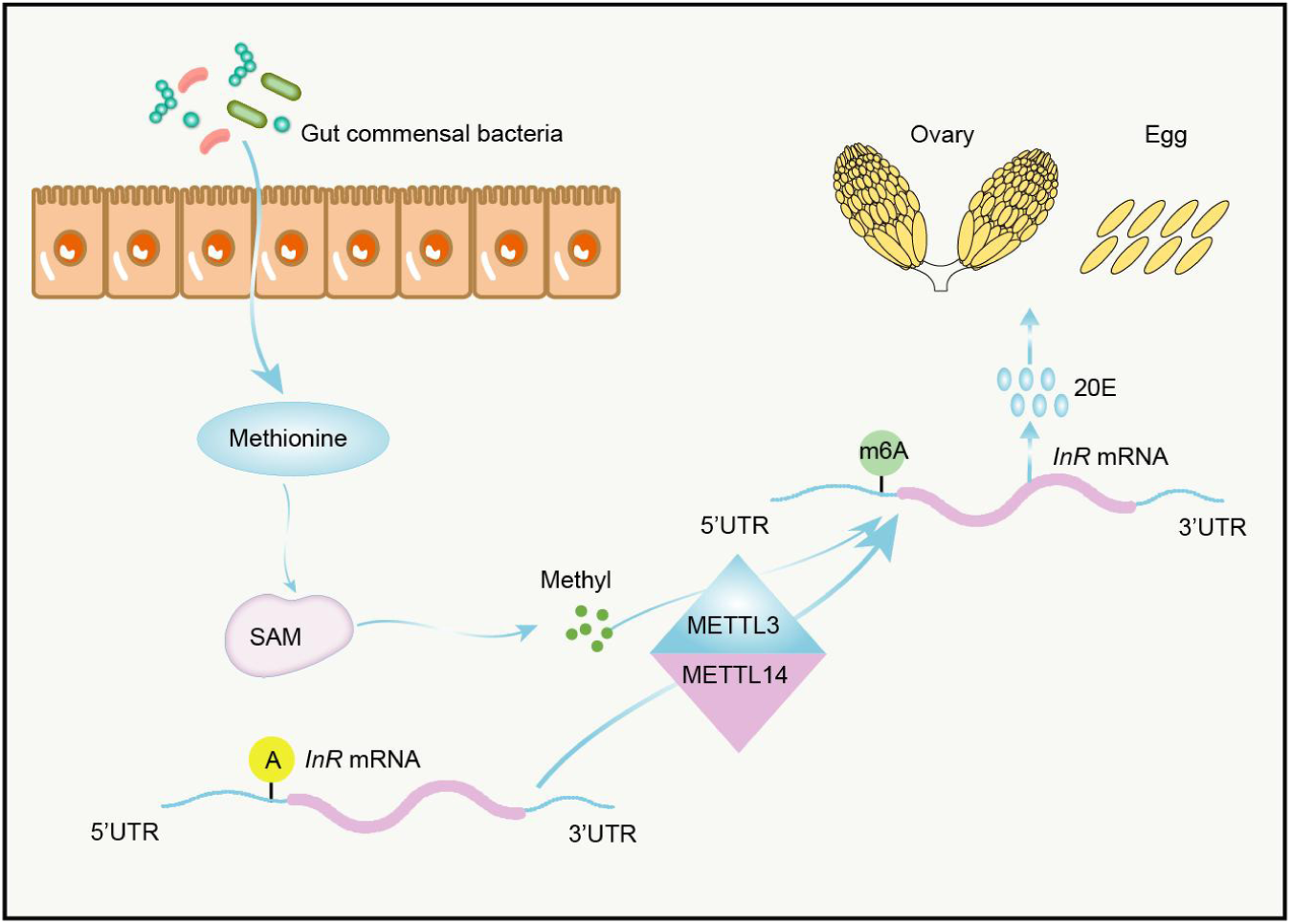
Schematic overview of how the gut commensal bacteria-derived methionine controls host reproduction by modulating RNA m6A methylation of *InR* in *B. dorsalis*. *E. hormaechei*-derived Met is converted to SAM, which provides methyl groups for the m6A modification of the target gene *InR* mRNA at its 5’ UTR region, thereby maintaining its stability. The resulting increased InR levels regulate ovarian development and fecundity by promoting ecdysone production.

It has been recently demonstrated that the insect gut microbiota ameliorates or complements the diet by providing vitamins, cholesterol, and amino acids, and influences multiple host traits, such as promoting larval growth ^32^ and host behavior ^33^. Previous studies have identified that the gut microbiota affects the oogenesis in *Drosophila* and that removal of *D. melanogaster* gut microbiota represses the oogenesis ^5^. However, our study is the first to decipher the mechanism by which the gut microbiota promotes host ovarian development and fecundity. We found that gut microbiota contributes to ovarian development by providing the essential amino acid methionine. Among the microbiota, *E. hormaechei* exhibited the most prominent promoting effect on the reproduction of female *B. dorsalis*. Previous research has highlighted the critical role of *E. hormaechei* in larval growth in host insects such as *D. melanogaster* ^34^ and *Musca domestica* ^35^, as well as its involvement in the detoxification processes of the housefly under exposure to heavy metal pollution ^36^. We previously identified a role of Enterobacter species, which are major components of the gut microbiota in *B. dorsalis* ^37^. Here, we reveal the critical role of *E. hormaechei* in insect reproduction through methionine production. Strikingly, methionine supplementation was sufficient to restore ovarian development and fecundity in germ free insects.

Nutrients are critical for successful female reproduction ^38^. For most insects, ovarian development is initiated by amino acid or protein intake ^39^. A diet lacking essential amino acids (i.e. histidine, isoleucine, or valine) reduced oviposition rates in *Drosophila* ^6^. Here, we reveal that methionine is key for ovarian development and fecundity in *B. dorsalis*. Methionine is an important essential amino acid responsible for several biological processes in animals ^40, 41^. In *Drosophila*, methionine is closely related to insect health and aging, and methionine restriction improves *Drosophila* health and extends its lifespan ^42^. Our study adds a new role for methionine in animal development.

Strikingly, we found that knockdown of the genes encoding the methyltransferase METTL3 and METTL14 blocked the promoting effect of methionine on ovarian development and fecundity, revealing that m6A is the major modulator of this process. In fact, Met can be converted to SAM under the action of S-adenosyl methioninesynthetase (MAT) ^29^, and then SAM provides methyl groups for mRNA methylation ^28^. We observed that methionine promoted the expression of methyltransferases METTL3 and METTL14 that catalyze RNA m6A methylation. Previous studies have shown that METTL3 and METTL14 are regulated by methionine levels ^13, 43^, which was also observed in this study. These findings suggest that methionine is critical for RNA m6A methylation levels in the ovary of *B. dorsalis* by providing the substrate to be converted to SAM, the donor of RNA m6A modification.

Our findings may not be restricted to insects since several reports have also demonstrated the great impacts of gut microbiota and amino acid metabolites on the reproductive health in humans ^7^. As observed in our study, methionine could regulate the expression of methyltransferase METTL3 in kidney cells of a polycystic kidney disease mouse model ^13^. Methionine deficiency was found to improve antitumor immunity by changing the m6A methylation of immune checkpoint transcripts in mice tumor cells ^14^. A recent study reported that methionine promotes the overall RNA methylation levels in human cells *in vitro* ^40^. Omics analysis has demonstrated that specific RNA m6A modifications are induced by gut microbiota in mice ^44^, and gut microbiota dysbiosis induced by antibiotics influences the murine transcriptome and m6A epitranscriptome ^45^. Thus, the mechanism by which gut microbiota affects host physiology at the epigenetic level by modulating m6A might be conserved across a range of organisms from invertebrates to vertebrates.

Furthermore, we showed that RNA m6A methylation is critical in the reproduction of female *B. dorsalis* and identified *InR* as the target gene of m6A modification. RNA m6A methylation regulates various biological processes including gametogenesis and embryogenesis by modulating RNA expression and metabolism in animals ranging from mammals ^46^ to insects ^20^. Recent research has also demonstrated the involvement of m6A in ovarian development of mice ^47^ and *Drosophila* ^48^. Thus, stabilization of mRNA transcripts by the m6A methylation machinery appears critical for ovarian development in various animal species, suggesting its conservation throughout evolution. We demonstrated that m6A modification of the target gene *InR* mRNA occurs at its 5’UTR region, thereby maintaining the mRNA stability of *InR* and regulating ovarian development and fecundity by promoting ecdysone production. The insulin pathway, which functions conservatively among animals ^49^, plays an important role as a nutritional sensor in insect reproduction, either independently or in combination with 20E or JH signaling ^38^. Our data reveal that m6A methylation of the transcript of *InR*, a master regulator of nutrient sensing, is critical for ovary maturation.

Based on the above results, we here propose a new model in which gut commensal bacteria-derived methionine regulates host reproduction by modulating RNA m6A methylation of the insulin receptor (Fig. 6). The gut commensal bacteria *E. hormaechei*-derived Met synthesizes SAM, providing methyl groups for the m6A reaction to increase *InR* m6A modifications, and then controlling *InR* expression in the ovary to activate ovarian development in *B. dorsalis*. These findings greatly improve our understanding of the complex networks that regulate female reproduction. To our knowledge, we have elucidated a previously unrecognized mechanism underlying how gut microbiota regulates host reproduction at the epigenetic level. Ultimately, these findings could suggest strategies for pest control and treating reproductive pathologies by targeting *InR* and the gut microbiota.

## Materials and methods

### Rearing of insects

Oriental fruit flies (*B. dorsalis*) were raised at the Institute of Horticultural and Urban Entomology, Huazhong Agricultural University (Wuhan, China), in climate-controlled chambers with a 27 ± 1°C temperature, 70 ± 5% RH (relative humidity), and 12: 12 h (L: D) photoperiod. The adults were fed with an artificial diet comprising 7.5% sugar, 2.5% honey, 2.5% yeast extract, and 87.5% water.

### Antibiotic treatment

For depletion of the gut microbiota, newly emerged female adults were fed with a liquid diet containing 3 mg/mL penicillin and 5 mg/mL streptomycin. The cotton pads were replaced every 24 h^1^.

### Bacteria quantification

The level of antibiotic elimination of gut microbes and colonization level of ABX flies were determined by colony-forming units from dissected guts^1^. The dissected guts were ground with sterile grinding pestles. The powder was then coated onto solid medium for 24 h of culture. Then, the colony-forming units (CFU) were counted.

### Gut bacterial culture, isolation, and identification

To isolate culturable bacteria from the guts of conventionally reared (CONV) females, the whole gut of eight females (4 days post eclosion) was first dissected and then collected with a sterile centrifuge tube, followed by the addition of 100 μL sterile water. Then, the gut was ground with sterile steel balls. A 100 ul diluent was then applied to an LB plate overnight. Individual genomic bacterial DNA was extracted from LB plates with an E.Z.N.A. Soil DNA kit (Omega, Norcross, GA, USA). The 16S rRNA gene was RT-PCR amplified to identify each bacterial isolate. The PCR conditions were 95 °C for 3 min; 35 cycles of 95 °C for 10 s, 55 °C for 10 s, and 72 °C for 20 s; 72 °C for 5 min. The primers were 27F and 1492R ^1^. Electrophoresis was then carried out on a 1% agarose gel to confirm the PCR products, followed by sequencing of the target PCR product. BLAST searching of the 16S rRNA sequence against the NCBI database was conducted to identify the culturable bacterial strains.

To isolate culturable bacteria in the fermentation of CONV female gut bacteria, the ground gut dilution was added to 100 ml LB liquid medium, followed by culturing overnight at 37 °C under shaking at 200 rpm. A 100 μL diluent was then applied to the LB plate overnight. The colony composition in the medium was detected by the above method.

### Association experiment between bacteria and female oriental fruit flies

The total culturable bacteria or isolated strains were placed in liquid LB media for culturing under appropriate conditions. CONV fruit flies were treated with antibiotics for 3 d and then fed culturable bacteria, as previously mentioned.

The cultivation experiments were carried out in a 250 ml shaking flask, and the culturable single strains screened for colonization and culturable total bacteria were added into the 100 ml LB medium, followed by culturing at 37 °C with 200 rpm shaking overnight. The oriental fruit flies were fed with bacteria at a final concentration of OD600 of 5.

### *E. hormaechei* localization by FISH

The gut samples of the female *B. dorsalis* were dissected on ice-cold PBS, which were then fixed in hybridization buffers (0.1M phosphate buffer and 4% polyformaldehyde, pH 7.0–7.5). The samples were then dehydrated and embedded in paraffin, sliced, and finally dyed with the FISH method following the previously described protocol ^1^. *E. hormaechei* probe for in situ hybridization was labeled with Cy3 at the 5’ and 3’ end of the *E. hormaechei* 16s sequence probe (Table S4).

### Reproductive phenotypic analysis

To analyze ovarian maturation, *B. dorsalis* females under different treatments were anesthetized on ice and dissected in cold PBS for determination of different ovarian development stages. Ovary images were taken with a stereo microscope fitted with a Nikon D5100 digital camera (Nikon, Tokyo, Japan). The Ovarian development of *B. dorsalis* could be divided into five stages according to the maturation degreein reference to a previous study, including previtellogenic (I), vitellogenic deposition (II), expectant stage of mature eggs (III), peak oviposition (IV), and final oviposition (V) ^26^. The ovarian proportions were calculated by dividing the number of ovaries at a certain stage by the total number of all ovaries.

To measure the laid egg number in each group, one sexually mature female and three males were placed in a self-developed egg counting device ^50^. The laid egg number in each cup was counted at 1, 3, 5, 7, and 9 d post-mating, and finally the total laid egg number was calculated and recorded. For each group, eight to ten biological replicates were performed.

### Nontargeted metabolomics analysis of hemolymph in *B. dorsalis*

After 4 d of antibiotic treatment, the hemolymph samples of the antibiotic-treated and control groups were extracted, with 6 biological replicates per treatment. After determining that the fertility of female flies in the antibiotic treatment group significantly decreased, 100 μl hemolymph samples were mixed with 500 μl of 80% methanol, followed by vortex shock, ice bath for 5 min, and centrifugation at 15000 r/min at 4 ℃ for 10 min. Then, 100 μl supernatant was taken and diluted with LC-MS grade water to 60% methanol content, and placed in a centrifuge tube with a 0.22 μm filter membrane, followed by centrifugation at 15000 r/min at 4 ℃ for 10 min. The filtrate was collected and injected for analysis. Hemolymph and metabolites were determined by LC-MS. A partial least squares-discriminant analysis (PLS-DA) was performed, and group differences were tested by permutational multivariate ANOVA (PERMANOVA).

### Targeted metabolomics analysis of *E. hormaechei*-derived essential amino acids

The essential amino acids for the metabolism of *E. hormaechei* were determined by UHPLC−MS/MS. A single clone of each wild type and mutant *E. hormaechei* derivative was placed in a 1.5 mL-tube and cultivated in 1 mL LB medium at 37°C overnight with orbital shaking at 220 rpm. Subsequently, the fermentation broth was centrifuged for 5 min at 15000 r/min. The bacteria were inoculated in a 200-mL shake flask containing 100 mL of MS medium. The culture was then incubated overnight at 37°C under orbital shaking at 200 rpm. Then, 20 ml fermentation broth was taken and placed in a centrifuge tube with 0.22 mm filter membrane for 15000 r/min shaking at 4 °C for 10 min. Subsequently, a 1-mL aliquot of clear supernatant was taken and transferred into an autosampler vial to perform UHPLC−MS/MS analysis.

### Construction of *E. hormaechei* ΔMetA mutant and detection of derived methionine

To verify the function of methionine produced by *E. hormaechei*, the gene MetA (Gene ID, EH72530959) was deleted in *E. hormaechei* by specialized transduction as described previously ^51^. In brief, the suicide vector pDM4-ΔMetA carrying chloramphenicol resistance was established by fusing two PCR products obtained by chloramphenicol of genomic sequences upstream and downstream MetA from *E. hormaechei*. Then, the pDM4-ΔMetA-CamR plasmid was transformed into the cells *E. coli* S17λpir. Subsequently, the plasmids were transfected into the wild-type *E. hormaechei* by bacterial conjugation. *E. hormaechei* mutants were screened for chloramphenicol resistance on LB media agar plates with chloramphenicol (43 μg/ml CmR). To complete the allele exchange required to generate ΔMetA mutants, the identified positive monoclonal exchange strains were streaked onto LB medium with 10% sucrose, and finally verified by Sanger sequencing.

### Culture conditions

To compare the growth of *E. hormaechei* and ΔmetA *E. hormaechei*, both strains were cultivated in LB, MS, M9, and M9 media with methionine at 37°C overnight. All *E. hormaechei* cells were washed with sterile distilled water three times and the cell density was measured with an OD600 Microplate Reader (Nano-300, All-Sheng, China). LB medium (10 g/L tryptone, 10 g/L NaCl, and 5 g/L yeast extract); MS medium (20 g/L glucose, 16 g/L ammonium sulfate, 2 g/L yeast extract, 1 g/L Potassium dihydrogen phosphite, 2 g/L Sodium Thiosulfate, 0.2 mg/L Vb12, 1 mL/L salt solution); M9 medium (1 g/L urea, 3 g/L KH2PO4, 6.78 g/L Na2HPO4, 0.1 g/L MgSO4 ·7H2O, 0.5 g/L NaCl, 15 g/L Agar)

### Measurement of methionine and SAM in *B. dorsalis*

To measure the effect of intestinal microorganisms on the titers of Methionine and SAM in oriental fruit flies after experimental treatment, hemolymph levels of methionine and SAM were measured by an ELISA kit (Jiangsu Baolai Biotechnology Co., LTD.) following the provided protocol. Hemolymph was collected and placed into a sterile centrifuge tube, followed by centrifugation at 4 °C and 3000 g/min for 15 min. The supernatant was collected and the protein content was quantified in a fresh centrifuge tube using a BCA kit. Then, hemolymph samples with equal concentration of proteins were added to detect SAM content. Sample readings at 450nm OD were plotted on a standard curve obtained by using the kit’s SAM-BSA standard for quantification of the methionine and SAM concentration in the tested samples, which were then normalized to the exact protein mass for further comparative analysis.

### Amino acid supplementation experiment in *B. dorsalis*

To verify the effect of methionine and valine on the host, newly emerging oriental flies (day 0) were divided into 4 groups: 1) CONV, 2) ABX, 3) ABX supplemented with methionine (ABX+Met), and 4) ABX supplemented with valine (ABX+Val). The CONV groups were fed according to normal rearing as stated. For the ABX groups, the flies were fed with antibiotics for three days and then changed to a sterile diet. For the ABX+Met groups, flies were fed with antibiotics for three days and then fed a sterile diet containing with 200 mM Met. For the ABX+Val groups, the flies were fed with antibiotics for three days and then fed a sterile diet containing with 200 mM Val. The CONV group and ABX group were fed with sterile diet to serve as a negative control.

### Dot blot

Total RNA was extracted from ten adults for each biological replicate with the RNAiso Plus reagent (TaKaRa, Otsu, Shiga, Japan) and denatured at 95°C for 3 min. RNA was spotted onto the HybondTM-N+ membrane and crosslinked for 30 min.

Following a 2 hour incubation in TBST (0.05% Tween in PBS) with 5% milk at room temperature, the membrane was incubated in blocking buffer (TBST 5% milk) with m6A antibody (ABclone, Wuhan, China, A22411) at 4°C overnight. Subsequently, the HybondTM-N+ membrane (Biosharp, China) was washed for three times with TBST for 15 min, followed by incubation with a secondary antibody in a blocking buffer for 1 h at room temperature. The membranes were detected by the ECL High sensitive Substrate (Monad, China). The dot blots were detected using a Multi Chemiluminescent Substrate Imaging System.

### Quantitative analysis of the methylation level of RNA m6A

The relative methylation value of m6A RNA was determined by using the EpiQuik m6A RNA Methylation Quantitative Kit (Epigentek, Farmingdale, NY, USA). Here, the number of strip wells required for the experiment was determined beforehand. Then, 200 ng of total input RNA was added to the strip wells, followed by capture antibodies (CA), detection antibodies (DA), and enhancer antibodies (EA). Finally, the developer solution (DS) was added to cease the color reaction. Then, the relative value of m6A RNA methylation was calculated based on the OD values.

### Antibody preparation and western blot

The METTL3 and METTL14 fragment obtained by PCR were transformed into the vector pET32a(+) for constructing recombinant vector pET32a(+)-METTL3 and pET32a(+)-METTL14. The vectors were transformed into *E. coli* BL21 to express the IPTG-induced fusion protein. Then, the protein was abundantly expressed and purified. The purified protein was used to immunize normal rabbits and prepare polyclonal antibodies.

For Western blot analysis, the isolated ovary tissue from *B. dorsalis* females was lysed in a RIPA buffer (GBCBIO Technologies, Guangzhou, China) that contained protease inhibitor (MedChemExpress, NJ, USA) to prepare the protein sample. Then, we centrifuged the homogenate at 12, 000 rpm and 4°C for 20 min, followed by collection of the supernatant. A 10% SDS–PAGE gel was used to separate the proteins, which were then transferred onto a PolyVinyliDene Fluoride (PVDF) membrane. After being blocked in TBST (0.05% Tween in PBS) with 5% milk at room temperature for 2 h, the membrane was incubated in a blocking buffer (TBST 5% milk) with primary antibody at 4 °C overnight. Then, the PVDF membrane was washed three times for 15 min in TBST, followed by incubation with a secondary antibody (polyclonal antibodies as described above) in the blocking buffer at room temperature for 1 h. The detection of protein bands was conducted using a Multi Chemiluminescent Substrate Imaging System. β-tubulin was used as loading control for the Western blot and detected with mouse anti-β-tubulin (Frdbio, Wuhan, China). The relative expression of the target protein was calculated as the ratio of the gray value of the target protein band to the gray value of β-tubulin.

### RNAi

A specific double-stranded RNA (dsRNA) was synthesized using the T7 RiboMAX Express RNAi System kit (Promega, Madison, WI, USA), and the primer sequence was listed in Table S4. *B. dorsalis* adults at 3 d after emergence were first anesthetized on ice, followed by microinjection of dsRNA (2 μg/μl) into the *B. dorsalis* ventral abdomen with a PV820 microinjector (WPI, USA). A negative control was constructed using equal amounts of dsEGFP.

### RNA extraction and qRT-PCR

Total RNA was extracted with the RNAiso Plus kit (TaKaRa, Otsu, Shiga, Japan). Then, the PrimeScript 1st strand cDNA Synthesis Kit with gDNA Eraser was used to reverse transcribe 1 μg total RNA onto the first strand of cDNA in each pool for qPCR (TaKaRa). qRT-PCR was carried out using a 10 μL reaction volume consisting of 5 μL qPCR Mix (Low ROX) (Yeasen, Shanghai), 2 μL cDNA (diluted at 1: 10), 0.4 μL 10 mM of each primer, and 2.2 μL RNase-free water. The qPCR cycling conditions were 5 min at 95 °C for initial denaturation, followed by 40 cycles of 95 °C for 10 s and 60 °C for 30 s. qRT-PCR was performed on the ABI 7500 real-time PCR system (Applied Biosystems, Foster City, CA, USA). The relative gene expression was calculated using the 2^(-ΔΔCt) method, and normalized to α-tubulin expression. For each biological replicate, three technical replicates were performed. The qRT-PCR primers are presented in Table S4.

### m6A methylated RNA immunoprecipitation qPCR (m6A-RIP-qPCR)

The m6A immunoprecipitation (m6A-RIP) procedure was conducted as previously described ^12^. Briefly, the ovaries were homogenized in a solution of cold RIPA lysis buffer (Beyotime, Shanghai, China), followed by centrifugation at 12, 000 g and 4 °C for 10 min. Then, 10 µg anti-m6A polyclonal antibody (ABclone, Wuhan, China) or normal Rabbit IgG protein was pre-incubated with 50 µl protein A/G beads for 1 h at 4°C. Subsequently, the ovarian lysate supernatant was added to the prepared antibody-bead mixture, which was then incubated at 4 °C overnight. Then, total RNA was extracted from the resulting RNA-protein complexes. About 10% of fragmented RNA was reserved as input control, and m6A enrichment was detected by RT-qPCR. The primers for m6A-RIP-qPCR are presented in Table S4.

### RNA sequencing (RNA-seq) and MeRIP-m6A-Seq analysis

Total RNA was isolated from CONV, ABX, ABX+EH Oriental fruit flies using RNAiso Plus reagent (TaKaRa, Otsu, Shiga, Japan). RNA quality was assessed on a NanoDrop ND-1000 (NanoDrop, Wilmington, DE, USA) and the RNA integrity was assessed by Bioanalyzer 2100 (Agilent, CA, USA). The mRNAs were purified using oligo(dT) magnetic beads (Thermo Fisher, CA, USA) and then were fragmented into segments 100 nt long (NEBNext® Magnesium RNA Fragmentation Module, USA). The untreated input control fragments (Input samples) and mRNA enriched with m6A-specific antibody (IP samples) were sequenced using Illumina NovaSeqTM 6000 by Lc-Bio Biotech Ltd (Hangzhou, China). Then, CleanData of the IP sample and Input sample were referenced to the *B. dorsalis* genome. R package exomePeak was employed for peak calling and diff Peak analysis with default settings. Using the R package ANNOVAR genetic structure and cross annotations. DESeq2 software was used to selected differentially expressed transcripts and genes with q <0.05 and |FC| ≥ 1.5. GO enrichment and KEGG enrichment analyses were used to analyze differentially expressed mRNAs. The enrichment analysis of differential m6A peaks was analyzed by GO, KEGG and MOTIF using Homer software.

### 20E determination

First, 150 μL hemolymph was collected from adult flies and mixed with 80% methanol of equal volume. The 300 μL mixture was vortexed and centrifuged at 12, 000 g for 10 min. Then, the upper methanol layer was transferred to a new tube and completely dried by using nitrogen. Then, the sample was re-suspended in an enzyme immunoassay buffer and subjected to enzyme immunoassays using the 20-Hydroxyecdysone Enzyme Immunoassay kit (Cayman Chemical Co., Ann Arbor, MI, USA, A05120) for estimation of the 20E titer.

### Cell culture and dual-luciferase reporter assays

Embryonic cell line of *B. dorsali* (Bd-EB, Laboratory stocks, unpublished data) was cultured at 28 °C in TNM-FH medium containing 10% fetal bovine serum (FBS). The wild-type InR 5’UTR sequences were mutated by replacing the adenosine bases (A) with guanine (G) using primers (Table S4) and cloned into the pGL4.26 reporter plasmid (Promega) to construct the plasmids pGL4.26 -InR-wt/mut-luc-SV40. The plasmid contained a mini promoter with a designated promoter region conjugated to firefly luciferase. The METTL3 and METTL14 sequences were constructed into the pAC5.1b/V5/His expression vector (Invitrogen). The primers used to construct vectors are shown in Table S4. Pre-treated Bd-EB were cultured into a 24-well plate followed by co-transfection of a reference reporter pGL4.73 plasmid containing the hRluc reporter gene (200 ng), pGL4.26 -InR-wt/mut-luc-SV40 (600 ng), and pAC5.1b/V5/His-METTL3 and METTL14(200 ng, ratio 1: 1) using the Polyethylenimine Liner (PEI) MW40000 (rapid lysis) Kit (YEASEN, China). At 72 h after transfection, measurement of luciferase activity was conducted with a Dual-Luciferase reporter assay system (Promega, WI, USA), which was then normalized to the luciferase activity of Renilla.

### mRNA stability assay

The METTL3 and METTL14 sequences were amplified and cloned into the shNC plasmid (Laboratory stocks) to construct the plasmids shMETTL3 and shMETTL14 using primers listed in Table S4. Cells transfected with shNC, shMETTL3, and shMETTL14 were incubated with 5 μg/ml actinomycin D (Act-D, Sigma, U.S.A), achieving RNA stability in *B. dorsalis* cells. Cells were then collected at indicated time, followed by isolation of RNA for RT-qPCR.

### Statistical analysis

All data are presented as the mean ± SEM from at least three independent experiments. Tests between the two groups were performed with a two-tailed Student’s *t*-test. Statistical analyses were performed using GraphPad Prism 8.0 software. Multiple comparisons were carried out with one-way analysis of variance (ANOVA) with Tukey’s test in SPSS 23.0 software (P<0.05).

## Contribution

H. Z. and Q. Z. conceived and designed the project. Q.Z. performed experiments and analyzed data. Z.D., and J.Q. did the gut bacterial identification experiments. Z.L., and P.L. made the graphs. Q.Z., W.Z., H.Z., B.L., and A.M.H. wrote the manuscript draft. All authors discussed and analysed the data, and modified the manuscript.

## Acknowledgments

This study was supported by the National Natural Science Foundation of China (no. 32220103009), China Agriculture Research System of MOF and MARA (CARS-26) and Hubei Hongshan Laboratory. We thank Claudine Neyen (EPFL, Lausanne, Switzerland) for help in editing the article. We would also like to thank Lin Qiu (Hunan agricultural university, Changsha, China) provided pGL4.73 plasmid.

---

**Supplementary Figure 1:**
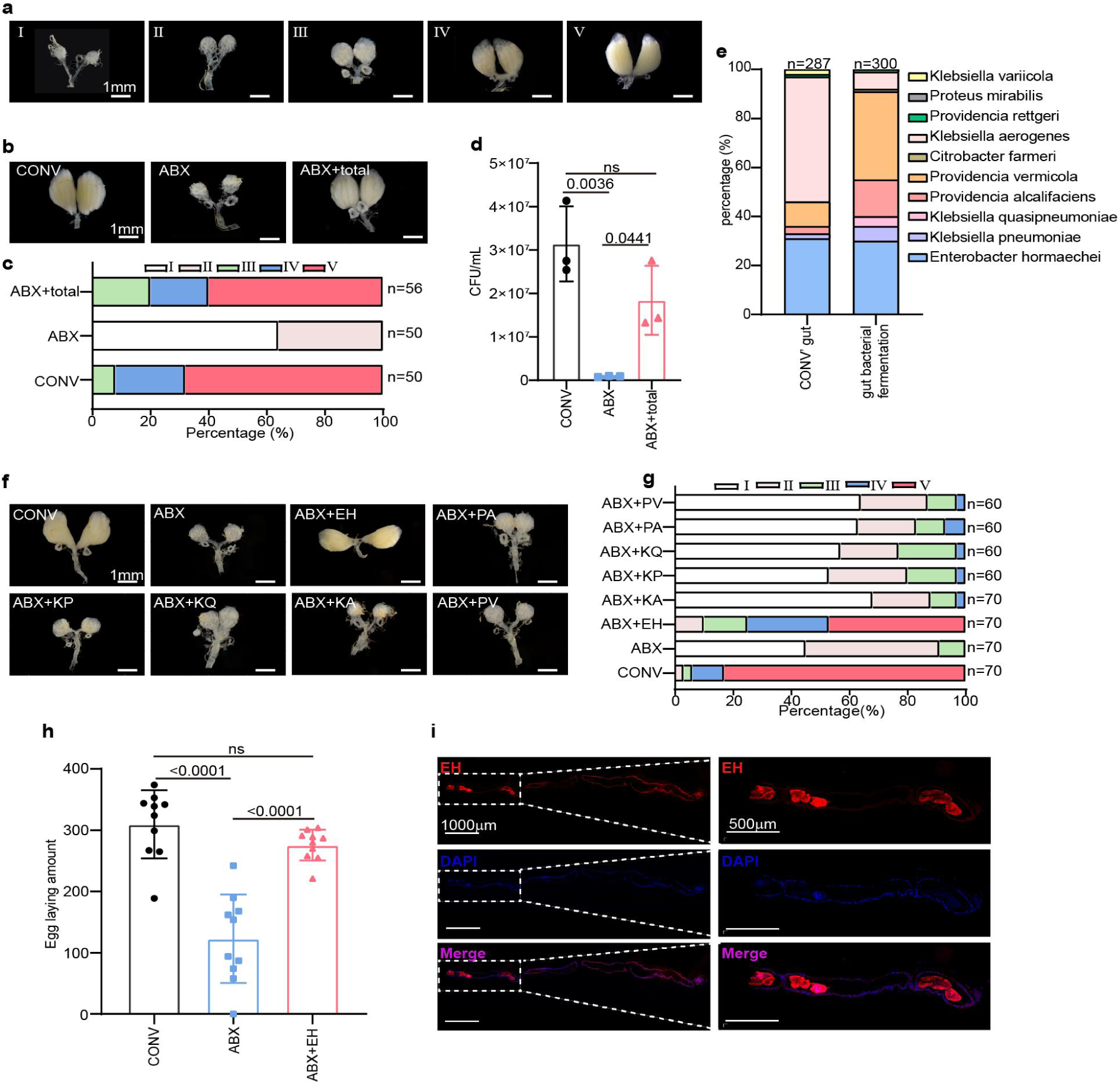
Gut bacteria promote ovarian development and fecundity in *B. dorsalis*. **a**, Representative morphology of *B. dorsalis* ovaries at five developmental stages. Stage I: The ovary is white and transparent, and the differentiation of the ovariole is not obvious; Stage II: The ovariole is visible and gradually expanded; Stage III: Oocytes increase rapidly, and yolk deposition increased; Stage IV: Some of the eggs mature; Stage V: Ovum maturation. **b**, **c**, Representative ovary morphology (b) and developmental stages (c) of ovaries analyzed in the ABX groups supplemented with total culturable bacteria. **d**, Total number of bacteria was determined by plating homogenized guts on LB plates and counting CFUs. **e**, Composition of gut bacteria in the gut and gut bacterial fermentation of CONV groups. **f**, **g**, Ovarian development of *B. dorsalis*, including representative ovary morphology (f) and development stages (g) of ovaries, mono-associated with *Enterobacter hormaechei* (EH), *Providencia alcalifaciens* (PA), *Klebsiella pneumoniae* (KP), *Klebsiella quasipneumoniae*, *Klebsiella aerogenes* (KA), *Providencia vermicola* (PV). **h**, Total number of eggs laid during oviposition. **i**, *E. hormaechei* localization in the gut was detected by fluorescence in situ hybridization (FISH). Red signals indicate *E. hormaechei* symbionts, whereas blue signals show host insect nuclei. Scale bars, 1000 µm and 500 µm. The number of independent biological replicates was three for **d**, while ten for **h.**

**Supplementary Figure 2:**
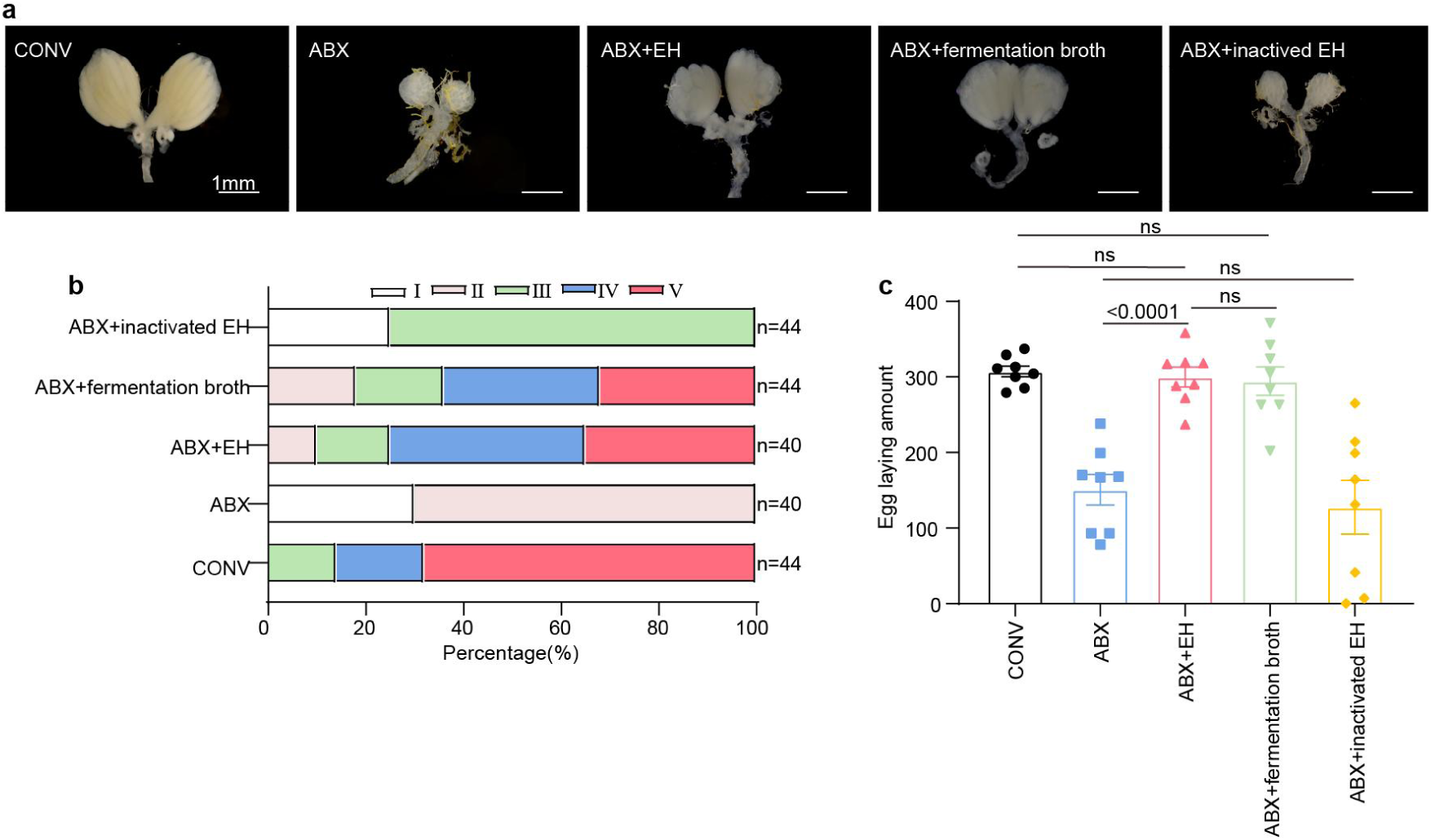
*E. hormaechei* fermentation controls *B. dorsalis* ovarian development and fecundity. **a-c** Representative ovary morphology (**a**) and developmental stages of ovaries (**b**) and total number of eggs laid during oviposition (**c**) were analyzed in the ABX group with different supplementation. CONV, conventional flies; ABX, antibiotic treated flies; ABX+ EH, ABX flies on a diet supplemented with *E. hormaechei*; ABX+fermentation broth, ABX flies on a diet supplemented with *E. hormaechei* fermentation broth; ABX+ inactived EH, ABX flies on a diet supplemented with heat-killed *E. hormaechei*.

**Supplementary Figure 3:**
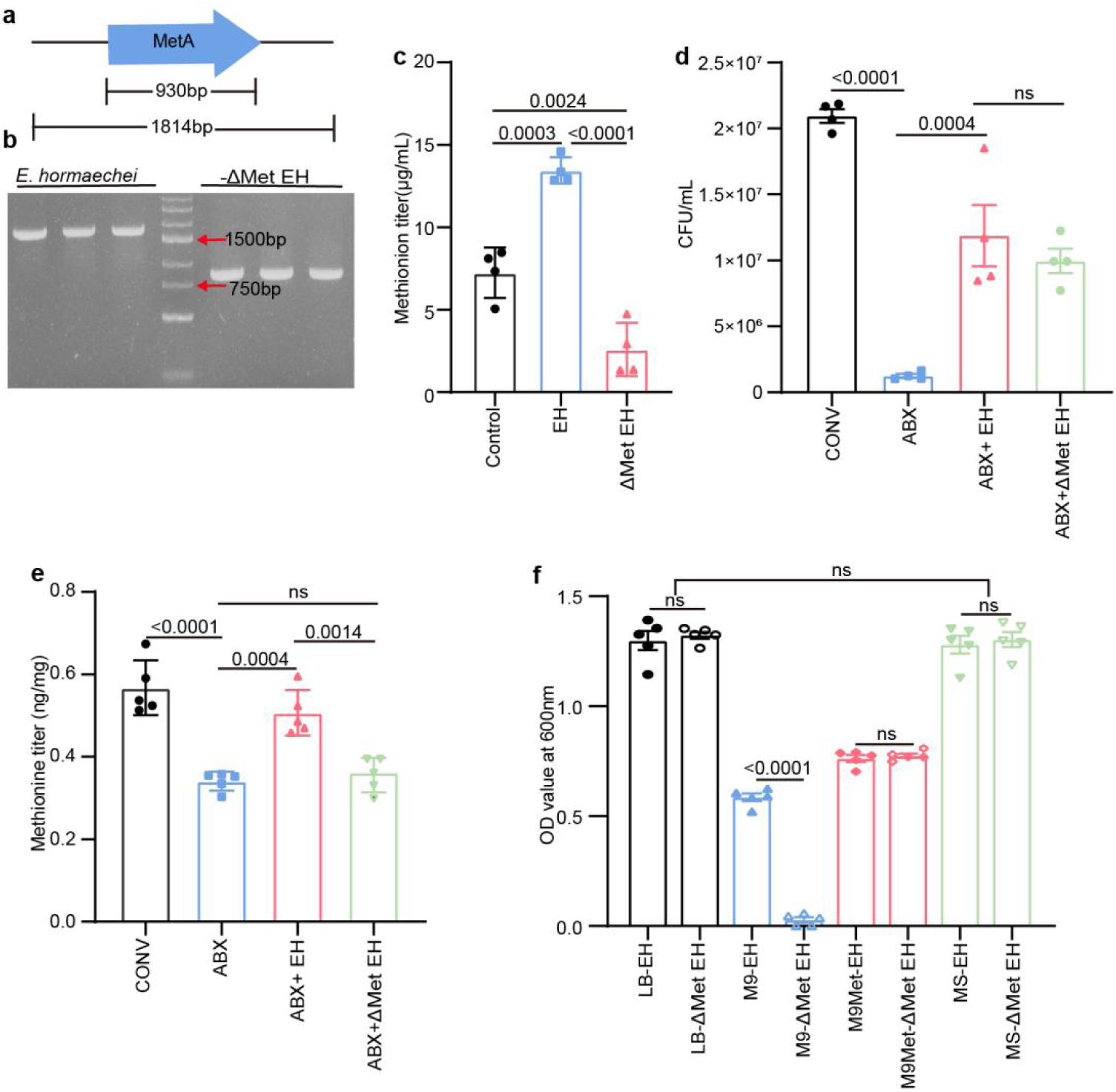
Functional study of *E. hormaechei* with methionine synthesis defect. **a**, Schematic diagram of MetA gene knockout in *E. hormaechei*. **b**, Agarose gel electrophoresis results of PCR products from WT and ΔMetA mutant *E. hormaechei*. **c**, Titer of methionine in WT and ΔMetA mutant *E. hormaechei*. **d**, Total number of bacteria was determined by plating homogenized guts on LB plates and counting CFUs. **e**, Titer of methionine in hemolymph of ABX flies mono-associated with WT or ΔMetA *E. hormaechei.* **f**, WT *and* ΔMetA *E. hormaechei* were grown overnight in LB, M9 medium, M9 medium containing methionine (M9Met), and MS medium at 37 °C and cell density was measured at OD600. The number of independent biological replicates was four for **d** and **e**, while five for **e** and **f.**

**Supplementary Figure 4:**
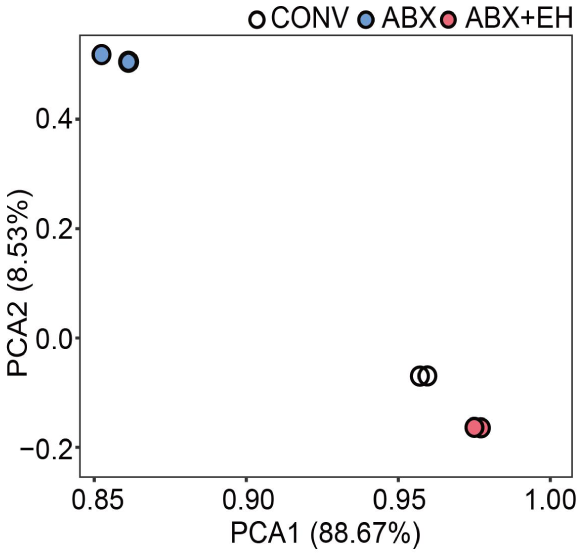
Principal component analysis (PCA) based on all differentially RNA m6A methylated modified mRNAs with the positions of samples in the space spanned. CONV, conventional flies; ABX, antibiotic treated flies; ABX+ EH, ABX flies on a diet supplemented with *E. hormaechei.* For each group, two independent biological replicates were conducted.

**Supplementary Figure 5:**
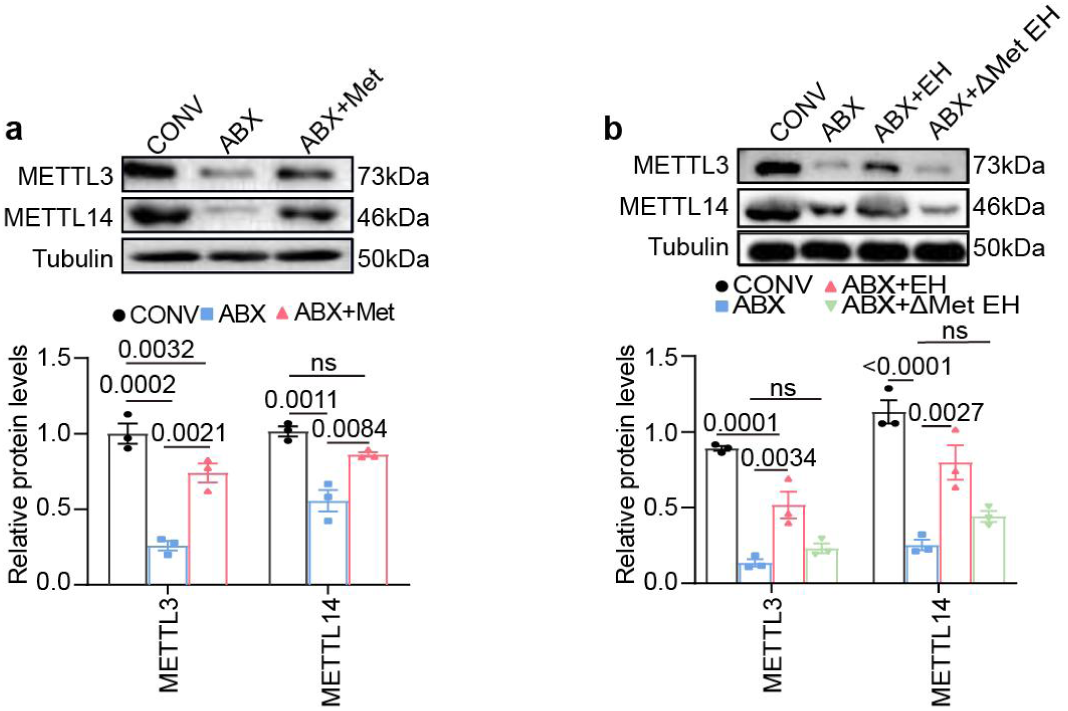
Methionine upregulates METTL3 and METTL14 expression. **a**, **b**, Relative expression of METTL3 and METTL14 determined by Western blot in the CONV, ABX, and ABX+Met groups (a) as well as in the CONV, ABX, ABX+WT *E. hormaechei*, and ABX+ ΔMetA *E. hormaechei* groups (b). Tubulin served as the loading control. The protein level was quantified by gray value and normalized to Tubulin. For each group, three independent biological replicates were conducted.

**Supplementary Figure 6:**
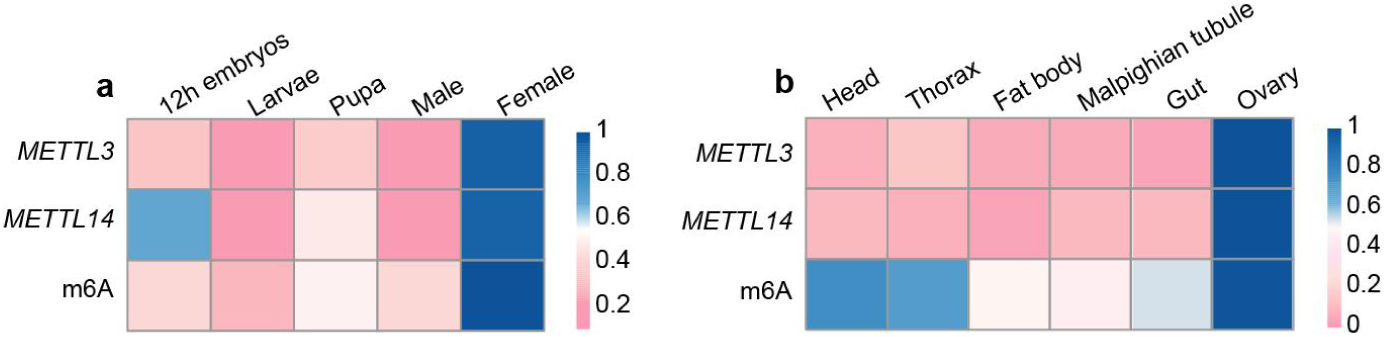
Spatiotemporal expression profiles of *METTL3*, *METTL14* and m6A. **a**, **b**, Heatmap showing the relative mRNA expression of *METTL3* and *METTL14* as well as m6A levels in different developmental stages (a) and tissues (b) in *B. dorsalis*. The number of independent biological replicates was three for **a** and **b**. The results were the mean of three replicates.

**Supplementary Figure 7:**
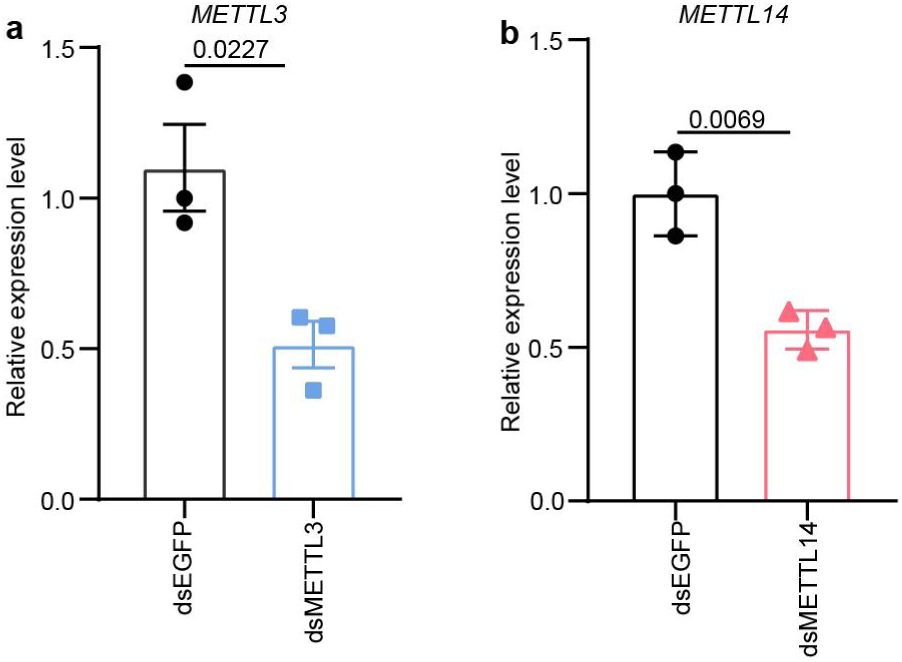
dsMETTL3 and dsMETTL14 efficiency detection in *B. dorsalis*. **a**, **b**, qRT-PCR quantifying the expression of METTL3 (a) and METTL14 (b) after injection of flies with dsEGFP, dsMETTL3, and dsMETTL14. The number of independent biological replicates was three for **a** and **b**.

**Supplementary Figure 8:**
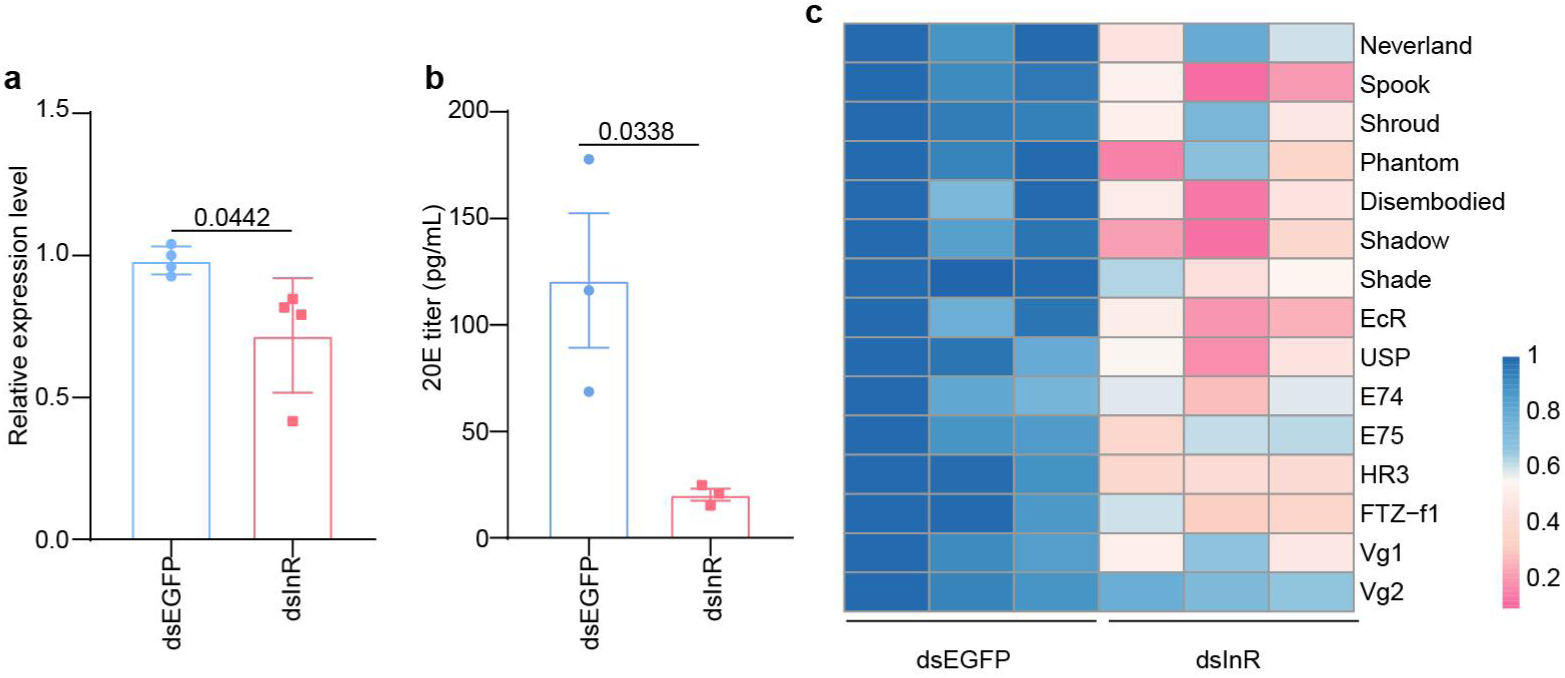
*InR* controls female reproduction by promotion of 20E biosynthesis in *B. dorsalis.* **a**, mRNA expression of *InR* in dsEGFP-injected and dsInR-injected female adults. **b**, 20E titer in the hemolymph of female adults injected with dsInR or dsEGFP at 48 h after post-injection by ELISA. **c**, Heat map showing the expression levels of ecdysone biosynthesis (*Neverland*, *Spook*, *Shroud*, *Phantom*, *Disembodied*, *Shadow*, *Shade*), 20E response (*EcR*, *USP*, *E74*, *E75*, *HR3*, *FTZ-f1*), and vitellogenin genes (*Vg1*, *Vg2*) in dsRNA-injected females by qRT-PCR. The number of independent biological replicates was four for **a**, while three for **b** and **c.**

## References

1. Yao Z, et al. Compartmentalized PGRP expression along the dipteran Bactrocera dorsalis gut forms a zone of protection for symbiotic bacteria. Cell Reports 41, (2022).

2. Yao Z, Wang A, Li Y, Cai Z, Lemaitre B, Zhang H. The dual oxidase gene BdDuox lregulates the intestinal bacterial community homeostasis of Bactrocera dorsias. The ISME Journal 10, 1037–1050 (2016).

3. Takeuchi T, et al. Gut microbial carbohydrate metabolism contributes to insulin resistance. Nature 621, 389–395 (2023).

4. O ’ Donnell MP, Fox BW, Chao P-H, Schroeder FC, Sengupta P. A neurotransmitter produced by gut bacteria modulates host sensory behaviour. Nature 583, 415–420 (2020).

5. Elgart M, Stern S, Salton O, Gnainsky Y, Heifetz Y, Soen Y. Impact of gut microbiota on the fly’s germ line. Nature Communications 7, (2016).

6. Leitão-Gonçalves R, et al. Commensal bacteria and essential amino acids control food choice behavior and reproduction. PLoS Biol15, e2000862 (2017).

7. Dai Z, Wu Z, Hang S, Zhu W, Wu G. Amino acid metabolism in intestinal bacteria and its potential implications for mammalian reproduction. Molecular Human Reproduction 21, 389–409 (2015).

8. Henriques SF, et al. Metabolic cross-feeding in imbalanced diets allows gut microbes to improve reproduction and alter host behaviour. Nat Commun 11, 4236 (2020).

9. Yang Y, Hsu PJ, Chen YS, Yang YG. Dynamic transcriptomic m6A decoration: writers, erasers, readers and functions in RNA metabolism. Cell Research 28, 616–624 (2018).

10. Patil DP, Pickering BF, Jaffrey SR. Reading m6A in the Transcriptome: m6A-Binding Proteins. Trends in Cell Biology 28, 113–127 (2018).

11. Meyer KD, Jaffrey SR. Rethinking m6A Readers, Writers, and Erasers. Annual Review of Cel and Developmental Biology 33, 319–342 (2017).

12. Ren Z, et al. MTA1 - mediated RNA m6A modification regulates autophagy and is required for infection of the rice blast fungus. New Phytologist2 35, 247–262 (2022).

13. Ramalingam H, et al. A methionine-Mettl3-N6-methyladenosine axis promotes polycystic kidney disease. Cel Metabolsm 33, 1234–1247.e1237 (2021)

14. Li T, et al. Methionine deficiency facilitates antitumour immunity by altering m(6)A methylation of immune checkpoint transcripts. Gut 72, 501–511 (2023).

15. Mu H, et al. METTL3-mediated mRNA N(6)-methyladenosine is required for oocyte and follicle development in mice. Cell Death Dis 12, 989 (2021).

16. Zhao BS, et al. m6A-dependent maternal mRNA clearance facilitates zebrafish maternal-to-zygotic transition. Nature 542, 475–478 (2017).

17. Haussmann IU, et al. m(6)A potentiates Sxl alternative pre-mRNA splicing for robust Drosophila sex determination. Nature 540, 301–304 (2016).

18. Lence T, et al. m(6)A modulates neuronal functions and sex determination in Drosophila. Nature 540, 242–247 (2016).

19. Yang X, et al. Epitranscriptomic regulation of insecticide resistance. Sci Adv 7, (2021).

20. Zhang GQ, et al. Dynamic FMR1 granule phase switch instructed by m6A modification contributes to maternal RNA decay. Nature Communications 13, (2022).

21. Clarke AR, et al. Invasive phytophagous pests arising through a recent tropical radiation: the Bactrocera dorsalis complex of fruit flies. Annu Rev Entomol 50, 293–319 (2005).

22. Guo Q, Yao Z, Cai Z, Bai S, Zhang H. Gut fungal community and its probiotic effect on Bactrocera dorsalis. Insect Science 29, 1145–1158 (2022).

23. Raza MF, et al. Gut microbiota promotes host resistance to low-temperature stress by stimulating its arginine and proline metabolism pathway in adult Bactrocera dorsalis. PLoS Pathog 16, e1008441 (2020).

24. Ren L, Ma Y, Xie M, Lu Y, Cheng D. Rectal bacteria produce sex pheromones in the male oriental fruit fly. Current Biology31, 2220–2226.e2224 (2021).

25. Cai Z, et al. Intestinal probiotics restore the ecological fitness decline of Bactrocera dorsalis by irradiation. Evolutionary Applications 11, 1946–1963 (2018)

26. Chou MY, Mau RF, Jang EB, Vargas RI, Piñero JC. Morphological features of the ovaries during oogenesis of the Oriental fruit fly, Bactrocera dorsalis, in relation to the physiological state. J Insect Sci 12, 1–12 (2012).

27. Weng H, et al. METTL14 Inhibits Hematopoietic Stem/Progenitor Differentiation and Promotes Leukemogenesis via mRNA m6A Modification. Cel Stem Cel 22, 191–205.e199 (2018).

28. Chiang PK, et al. S-Adenosylmethionine and methylation. Fasebj 10, 471–480 (1996).

29. Mentch SJ, Locasale JW. One-carbon metabolism and epigenetics: understanding the specificity. Annals of the New York Academy of Sciences 363, 91–98 (2015).

30. Liu J, et al. A METTL3-METTL14 complex mediates mammalian nuclear RNA N6-adenosine methylation. Nat Chem Biol 10, 93–95 (2014).

31. Wang P, Doxtader KA, Nam Y. Structural Basis for Cooperative Function of Mettl3 and Mettl14 Methyltransferases. Molecular Cell 63, 306–317 (2016).

32. Matos RC, et al. D-Alanylation of teichoic acids contributes to Lactobacilus plantarum-mediated Drosopliha growth during chronic undernutrition. Nature Microbiology 2, 1635–1647 (2017).

33. Kang WK, et al. Vitamin B12 produced by gut bacteria modulates cholinergic signalling. Nature Cell Biology 26, 72–85 (2024).

34. Kang r, Li y, Bai Q, Gao Y, Iiu W. Enterobacter hormaechei promotes the growth and development of Drosophila melanogaster. (2020)

35. Zhang Q, et al. Enterobacter hormaechei in the intestines of housefly larvae promotes host growth by inhibiting harmful intestinal bacteria. Parasites & Vectors 14, (2021).

36. Yin Y, et al. Antagonistic effect of the beneficial bacterium Enterobacter hormaechei against the heavy metal Cu2+ in housefly larvae. Ecotoxicology and Environmental Safety 272, (2024).

37. Wang HX, Jin L, Peng T, Zhang HY, Chen QL, Hua YJ. Identification of cultivable Lsaroa Bectc bacteria in the intestinal tract of radis from three different populations and determination of their attractive potential. Pest Management Science70, 80–87 (2014).

38. Roy S, Saha TT, Zou Z, Raikhel AS. Regulatory Pathways Controlling Female Insect Reproduction. Annu Rev Entomol 63, 489–511 (2018).

39. Hansen IA, Attardo GM, Park JH, Peng Q, Raikhel AS. Target of rapamycin-mediated amino acid signaling in mosquito anautogeny. Proc Natl Acad Sci USA 101, 10626–10631 (2004).

40. Liu X, et al. Gut microbial methionine impacts circadian clock gene expression and reactive oxygen species level in host gastrointestinal tract. Protein Cell 14, 309–313 (2023).

41. Gu X, et al. SAMTOR is an S-adenosylmethionine sensor for the mTORC1 pathway. Science 358, 813–818 (2017).

42. Parkhitko AA, et al. A genetic model of methionine restriction extends Drosophila health- and lifespan. Proceedings of the National Academy of Sciences of the United States of America 118, (2021).

43. Villa E, et al. mTORC1 stimulates cell growth through SAM synthesis and m6A mRNA-dependent control of protein synthesis. Molecular Cell 81, 2076–2093.e2079 (2021)

44. Jabs S, et al. Impact of the gut microbiota on the m6A epitranscriptome of mouse cecum and liver. Nature Communications 11, (2020).

45. Yang M, et al. Antibiotic - Induced Gut Microbiota Dysbiosis Modulates Host Transcriptome and m6A Epitranscriptome via Bile Acid Metabolism. Advanced Science, (2024).

46. Yao H, et al. scm6A-seq reveals single-cell landscapes of the dynamic m6A during oocyte maturation and early embryonic development. Nature Communications 14, (2023).

47. Sun X, Zhang J, Jia Y, Shen W, Cao H. Characterization of m6A in mouse ovary and testis. Clin Transl Med 10, e141 (2020).

48. Hongay CF, Orr-Weaver TL. Drosophila Inducer of MEiosis 4 (IME4) is required for Notch signaling during oogenesis. Proc Natl Acad Sci USA 108, 14855–14860 (2011).

49. Abrisqueta M, Süren-Castillo S, Maestro JL. Insulin receptor-mediated nutritional signalling regulates juvenile hormone biosynthesis and vitellogenin production in the German cockroach. Insect Biochemistry and Molecular Biology 49, 14–23 (2014).

50. Li T, Ye Y, Wu P, Luo R, Zhang H, Zheng W. Proteasome β3 subunit (PSMB3) controls female reproduction by promoting ecdysteroidogenesis during sexual maturation in Bactrocera dorsalis. Insect Biochemistry and Molecular Biology 157, (2023).

51. Khider M, Willassen NP, Hansen H. The alternative sigma factor RpoQ regulates colony morphology, biofilm formation and motility in the fish pathogen Alivibrio salmonicida. BMC Microbiology 18, (2018).

