## Supplemental Table 1 for "Gut Commensal Bacteria-Derived Methionine is Required for Host Reproduction by Modulating RNA m6A Methylation of the Insulin Receptor"

Table S1 Composition of gut bacteria in CONV's gut and gut bacterial fermentation.

| **Representative bacterial strain** | **CONV'S gut** | **gut bacteria fermentation** |
|---|---|---|
| *Enterobacter hormaechei* | 87（30. 31%） | 91（30.43%） |
| *Providencia vermicola* | 28（9. 76%） | 108（36.12%） |
| *Providencia alcalifaciens* | 10（3. 48%） | 45（15.05%） |
| *Klebsiella aerogenes* | 146（50. 87%） | 20（6.69%） |
| *Klebsiella pneumoniae* | 5（1. 74%） | 18（6.02%） |
| *Klebsiella quasipneumoniae* | 1（0. 35%） | 12（4.01%） |
| *Providencia rettgeri* | 4（1. 39%） | 3（1%） |
| *Citrobacter farmeri* | 0（0%） | 2（0.67%） |
| *Klebsiella variicola* | 5（1.74%） | 0（0%） |
| *Proteus mirabilis* | 1（0. 35%） | 0（0%） |
| *Sum of colonies* | 287 | 300 |
