## Supplemental Table 2 for "Gut Commensal Bacteria-Derived Methionine is Required for Host Reproduction by Modulating RNA m6A Methylation of the Insulin Receptor"

Table S2 Extracellular metabolites in *E. hormaechei*

| (μg/mL) | Val | Thr | Ile | Leu | Lys | Met | His | Phe | Arg | Trp |
|---|---|---|---|---|---|---|---|---|---|---|
| | 30.69 | 0.09 | 14.69 | 9.46 | 0.00 | 3.94 | 0.00 | 0.20 | 0.58 | 0.26 |
| *E.hormaechei* | 23.66 | 0.09 | 13.64 | 9.02 | 0.00 | 5.19 | 0.40 | 0.28 | 0.00 | 0.26 |
| | 37.96 | 0.12 | 16.00 | 10.57 | 0.00 | 6.15 | 0.00 | 0.36 | 0.69 | 0.35 |
